# Cancer cell population growth kinetics at low densities deviate from the exponential growth model and suggest an Allee effect

**DOI:** 10.1101/585216

**Authors:** Kaitlyn E. Johnson, Grant Howard, William Mo, Michael K. Strasser, Ernesto A. B. F. Lima, Sui Huang, Amy Brock

## Abstract

Models of cancer cell population expansion assume exponential growth kinetics at low cell densities, with deviations from exponential growth only at higher densities due to limited resources such as space and nutrients. However, recent pre-clinical and clinical observations of tumor initiation or recurrence indicate the presence of tumor growth kinetics in which growth rates scale with cell numbers. These observations are analogous to the cooperative behavior of species in an ecosystem described by the ecological principle of the Allee effect. In preclinical and clinical models however, tumor growth data is limited by the lower limit of detection (i.e. a measurable lesion) and confounding variables, such as tumor microenvironment and immune responses may cause and mask deviations from exponential growth models. In this work, we present alternative growth models to investigate the presence of an Allee effect in cancer cells seeded at low cell densities in a controlled *in vitro* setting. We propose a stochastic modeling framework to consider the small number of cells in this low-density regime and use the moment approach for stochastic parameter estimation to calibrate the stochastic growth trajectories. We validate the framework on simulated data and apply this approach to longitudinal cell proliferation data of BT-474 luminal B breast cancer cells. We find that cell population growth kinetics are best described by a model structure that considers the Allee effect, in that the birth rate of tumor cells depends on cell number. This indicates a potentially critical role of cooperative behavior among tumor cells at low cell densities with relevance to early stage growth patterns of emerging tumors and relapse.

**Author Summary:** The growth kinetics of cancer cells at very low cell densities are of utmost clinical importance as the ability of a small number of newly transformed or surviving cells to grow exponentially and thus, to “take off” underlies tumor formation and relapse after treatment. Mathematical models of stochastic tumor cell growth typically assume a stochastic birth-death process of cells impacted by limited nutrients and space when cells reach high density, resulting in the widely accepted logistic growth model. Here we present an in-depth investigation of alternate growth models adopted from ecology to describe potential deviations from a simple cell autonomous birth-death model at low cell densities. We show that our stochastic modeling framework is robust and can be used to identify the underlying structure of stochastic growth trajectories from both simulated and experimental data taken from a controlled *in vitro* setting in which we can capture data from the relevant low cell density regime. This work suggests that the assumption of cell autonomous proliferation via a constant exponential growth rate at low cell densities may not be appropriate for all cancer cell growth dynamics. Consideration of cooperative behavior amongst tumor cells in this regime is critical for elucidating strategies for controlling tumor cell growth.

## Introduction

The classical formulation of tumor growth models begins with the assumption that early stage tumor growth dynamics are driven by cell-autonomous cell proliferation, manifested as an exponential increase in cell number, where the growth rate is proportional to the number of cells present and is captured by a single growth rate constant. However, the ability to measure early stage tumor growth, at a stage when only low cell numbers are present, is not feasible in the clinic, as patients present only after the tumor has exceeded the lower limit of detection (about 1 million cells on a typical CT scan(1)). Recent findings in preclinical mouse models (2) and from clinical outcomes following tumor resection (3) reveal that tumor growth at low tumor cell densities does not match the expectation of exponential growth. These findings give rise to an intriguing possibility: does tumor cell growth deviate from the model of exponential growth at low tumor cell densities? In this study, we ask whether early stage tumor growth kinetics exhibits a behavior analogous to a principle in ecology known as the Allee effect, in which the fitness of a population, measured by the per capita growth rate, scales with population size at low population sizes. In ecology, the Allee effect arises due to cooperative growth, such as cooperative predation, feeding, and mating systems (4). In tumors, subclonal interactions are observed among cells, with specific subpopulations releasing signaling molecules critical to the growth of other subsets of cells (5–9). Thus cancer cell growth may exhibit cooperative interactions analogous to those amongst species in an ecosystem.

The ability to describe and predict tumor growth is essential to developing strategies to eradicate cancer cell populations (10,11). Understanding tumor growth kinetics at low cell numbers is of clinical importance as they govern tumor initiation, treatment response, and recurrence. In ecology, the Allee effect has informed strategies for the control of invasive species (12), and applying ecological principles to control tumor growth is a growing interest (13–22). A better understanding of the factors that govern tumor cell growth at early stages could guide the manipulation of subpopulation dynamics to disrupt cooperative growth as a therapeutic strategy for preventing metastasis and tumor progression.

The assumption that early stage tumor growth kinetics matches exponential growth models is widely accepted in the field, with much of the focus of tumor growth modeling on how best to describe the slowing of growth rate at larger tumor sizes (16,23–26). The most common model used to describe a wide variety of solid tumors is the logistic or related growth models (16,23,24), in which tumors are assumed to grow (near) exponentially from a single cell until the tumor size approaches carrying capacity, at which time tumor growth slows and approaches net-zero (a plateau) as cancer cells are met with limited resources of space and nutrients. A model more consistent with realistic faster flattening of the growth curve towards the carrying capacity is the Gompertzian growth model (23,26). Such models are convenient because they not only match clinically observed growth kinetics, but can be observed in an *in vitro* setting simply by plating cells in a dish and allowing them to grow until they approach confluence, at which point limited space and nutrients mimics the carrying capacity observed *in vivo*. Deviations from early exponential growth have also been modeled, but in a phenomenological manner, assuming non-constant growth behaviors(27,28).

A mechanistic view for deviations from (near) exponential early growth is the Allee effect which results from cell-cell interactions in the low-cell number regime which can promote growth. Indeed, recently deviations from exponential growth in early stages of tumor growth have been observed in glioblastoma, where patient brain tumors were resected and monitored over time for relapse (3). These studies of relapsed tumor growth revealed that the observed growth rate at the clinically detectable stages of tumor growth failed to match models of simple logistic growth, and instead were better described by a weak Allee effect model. In the weak version of the Allee effect, populations grow at a much slower rate at very low tumor cell numbers, but continue to grow for any initial population size. By contrast, a strong Allee effect describes a population that becomes extinct below a threshold initial population size. In ecology, both strong and weak Allee effects are observed (4). While the observation of a weak Allee effect in glioblastoma recurrence is certainly provocative, it is limited by the fact that the earliest stages of tumor growth from low cell densities cannot be easily detected *in vivo*, and thus the critical measurements at the relevant regime cannot be captured with current imaging technologies. While numerous studies have investigated the manifestation of the Allee effect in ecology (29–32) and a few have posed theoretical implications of the Allee effect of cooperative kinetics in cancer growth (33–37), none have performed an indepth quantitative analysis of cancer cell proliferation kinetics captured in the low cell density regime to assess the relevance of growth models that incorporate an Allee effect.

In this study, we investigate the behavior of various structurally distinct models of tumor growth representing alternative hypotheses of growth dynamics that consider the Allee effect. We present a framework for the analysis of cancer cell growth at low cell densities in a controlled *in vitro* setting. Monitoring growth *in vitro* allows for studying the effects of cell number on growth in the absence of confounding factors, such as the immune system interactions and tissue microenvironmental factors, in order to test explicitly the dependence of growth dynamics on cell density. We take advantage of recent technological advances that allow for the seeding of a precise small initial cell number and the ability to measure cell number at single cell-resolution and at high temporal resolution in order to capture accurate growth kinetics in the low cell density regime where the Allee effect would be most relevant and which cannot be studied *in vivo*. We focus our examination in the low cell density regime (<200 cells in a 1 mm^3^ well) and therefore our modeling analysis excludes additional terms that describe the slowing of cancer cell population growth at higher densities where competition for limited resources becomes relevant. We examine the average behavior of three models of increasing complexity: the exponential growth model, a strong Allee model, and an extended Allee model that can be either strong or weak. To account for the inherent stochasticity in population size revealed at very small population sizes, we develop seven stochastic models whose average behavior follows one of the deterministic models, with each stochastic model representing a different hypothesis of the mechanism underlying the growth kinetics. For each stochastic model, we perform a parameter estimation using the method of moments (25) and use model selection to identify the model most likely to describe the growth data(38). This is performed for both a simulated data set and the *in vitro* BT-474 data to test the hypotheses that our framework reveals an alternative tumor growth model that includes an Allee effect.

## Results

### The deterministic strong and weak Allee effect models

This work studies stochastic growth models because of the inherent stochasticity of cell growth processes in the regime of small cell numbers that is our focus. However, the structure of the functional forms of the kinetic equations describing the cell number changes can be understood within a deterministic framework, thus providing a link to the historical and most widely implemented tumor growth models. At the core of our models are the following three deterministic phenomenological models of increasing complexity that describe cell population growth kinetics. The first model represents the “null model” of tumor growth (23) where the growth rate (*dN*/*dt*) is proportional to the number of cells present, *N*, and a single growth rate constant, g, resulting in the classical exponential growth model.

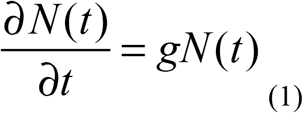

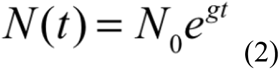

This model describes *N* cells that exhibit cell autonomous proliferation (Figure 1A) and a constant per capita growth rate((*dN*/*dt*)/N) given by the growth rate constant *g* over time and cell number (Figure 1B) for initial cell numbers N_0_ = 3, 8, and 16 cells displayed in Figure 1 A, B, & C. The normalized growth rate (log(N(t)/N_0_)), is constant for each initial condition, with all growth curves falling on a line of equal slope (Figure 1C). Equation 1 & 2 represent the well-known exponential growth model and the simplest of the tumor growth models analyzed.

**Figure 1.**
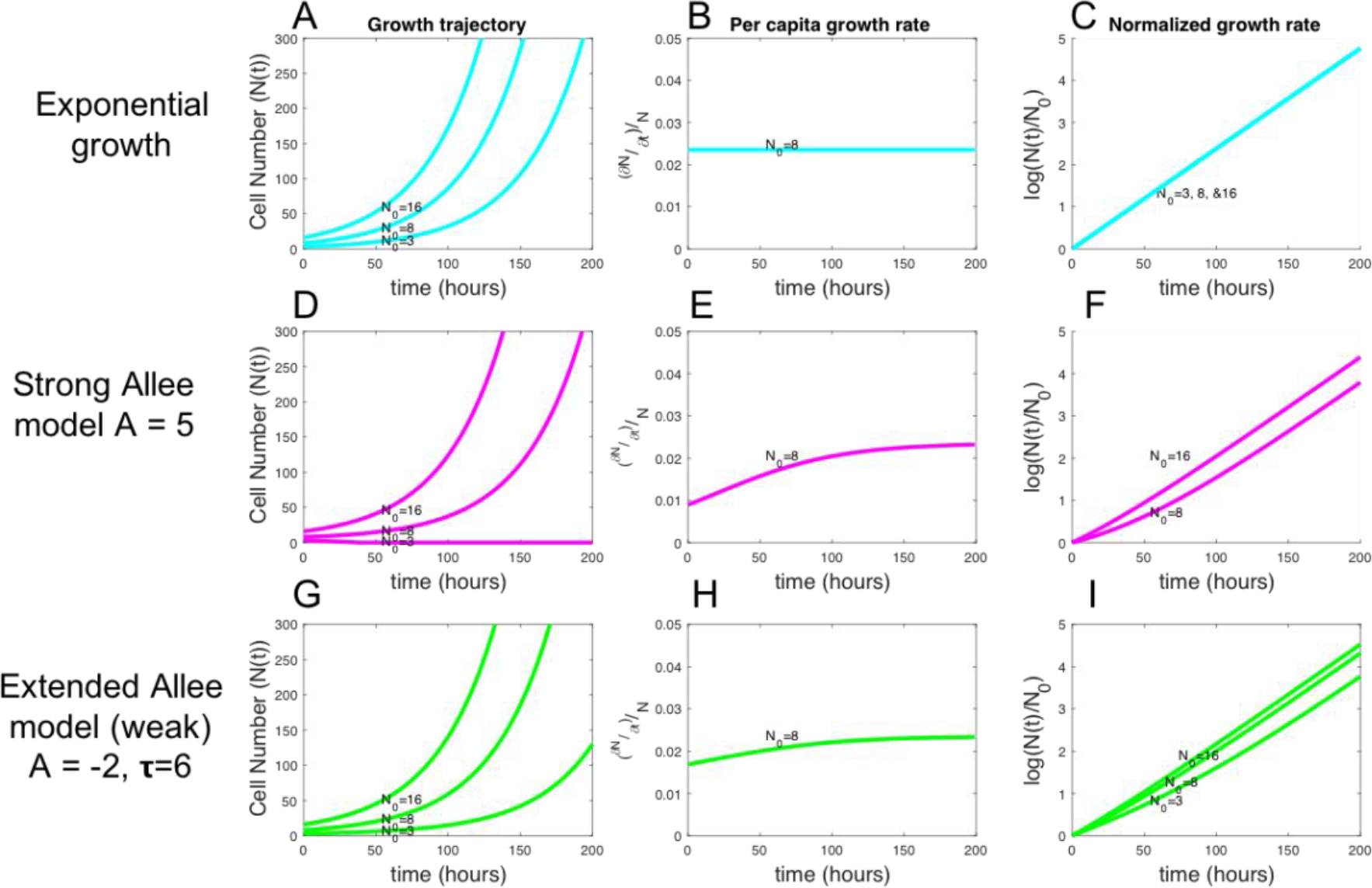
Average behavior of exponential, strong, and weak Allee models for different initial conditions. (a) (d) & (g) Deterministic growth curves of the exponential growth model (blue), the strong Allee model (pink), and the weak Allee model (green) respectively, shown for N_0_ = 3, 8, and 16 for all models. (b) (e) & (h) Per capita growth rates demonstrate that growth rate increases in time with cell number for both Allee models. (c) (f) & (i) For normalized cell numbers, a clear difference is observed in the slopes depending on the initial cell number for both Allee models.

Most departures from the exponential growth model of cancer cells (Eq.1 & 2) describe cancer cell growth where the growth rate is proportional to the number of cells present but with a growth rate that may depend explicitly on *N*. For example, in the classical formulation of the logistic growth model(23), the growth rate is characterized by a growth rate constant *g* multiplied by an additional carrying capacity (*K*) term dependent on *N* of (*1-N/K*), as shown below:

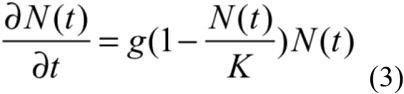

The logistic growth model describes cells in two regimes: when N≪K the *N/K* term is negligible and the cells essentially exhibit exponential growth, and when N is near K net growth rate (*dN/dt*) slows towards zero as N approaches K and the (*1-N/K)* term approaches zero. Much like this description of slowing of growth rate at high cell numbers, the second model, the strong Allee model, introduces an additional Allee effect term (*1-A/N*) that is dependent on N to the per capita growth rate:

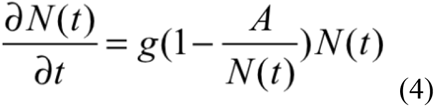

This model describes N cells whose net growth rate exists in two distinct regimes: when N is less than the Allee threshold (A), the Allee effect term (*1-A/N*) in Equation (4) becomes negative and the net growth rate *(dN/dt)* becomes negative, predicting the population will ultimately go extinct (Figure 1D, N_0_=3). When N(t) is near A but larger than A, the net growth rate is slowed by a factor of (*1-A/N*) (Eq. 4) but remains positive, resulting in a growth rate that scales with cell number, as can be seen for the per capita growth rate over time for N_0_=8 (Figure 1E). When N(t) is much larger than A, the Allee effect term (*1-A/N*) becomes negligible and the cell population begins to behave like in the exponential growth model (Figure 1D, E). This behavior in which a population is predicted to go extinct below a critical number describes a strong Allee effect. The expected scaling of the normalized growth rate (log(N(t)/N_0_)) demonstrates the expected differences in net growth rate based on initial seeding number for a strong Allee model (Figure 1F). As expected, an initial condition of N_0_<A results in a negative growth rate and therefore negatively sloped line starting at 0 (not shown). This model is able to explain the threshold-like behavior observed in pre-clinical studies of engrafted tumors in mouse (2); namely, below a threshold number of inoculated cells, tumors never form. To account for the weak Allee effect behavior, in which the growth rate is always greater than zero for any N_0_, we introduce the third deterministic model: the extended Allee model.

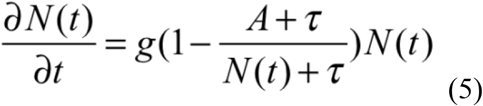

This model is similar to the strong Allee model (Eq.4) but introduces an additional parameter τ which allows the model to exhibit either a strong Allee effect when A is positive, or a weak Allee effect when τ >|A| and A<0. When weak Allee conditions hold, at low N the (*A+τ/N+τ*) term always remains less than 1, keeping the net growth rate positive but resulting in a growth rate that approaches zero as N decreases (although it never goes below zero to predict extinction). Figure 1G, H, I display the behavior of the extended Allee model with parameters that produce a weak Allee effect. The extended Allee model explains potential weak Allee effects, such as those observed in the glioblastoma resection case (3). See Table S1 for a complete description of each of the three deterministic models, their parameters, and their behaviors.

### Extension to stochastic growth models

Given that the growth kinetics are measured here in very small cell populations, where the expected variability of individual cell behavior with respect to division (‘birth’) and ‘death’ events (which jointly determine net growth rate *dN/dt*) is high, this scenario can give rise to apparent growth rates that deviate significantly from the average population behavior. Thus, the deterministic models may not reflect the actual behavior of individual cell number trajectories of each small cell-number initial population. To consider the inherent stochasticity in the birth-death process observable in our experimental system instead of fitting the averaged data to the deterministic models, we developed corresponding stochastic models whose expected cell numbers *N(t)* are equivalent to that predicted by the ODE of the deterministic models described above (Eqs. 1, 4, & 5). For each deterministic model structure, the total number of cells N(t) is modeled by the ODEs Eq. 1, 4, & 5 above. The time evolution of N(t) for the stochastic models are defined by the following birth and death events (Figure 2A):

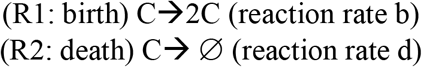

With propensities:

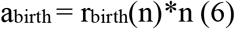

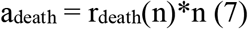

Where r_birth_(n) and r_death_(n) describe the rate at which the events occur, which may depend on the number of cells, n, present or be constant. The probability of an event i happening in an infinitesimal time step dt is given generally by:

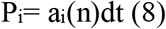

Where a_i_ describes the propensity for that event *i* to occur. In this model of tumor growth, the only possible events are birth or death events, and in all cases the propensity of an event to occur is a function of *n*, as it is a first order reaction described by the schematic in Figure 2A.

**Figure 2.**
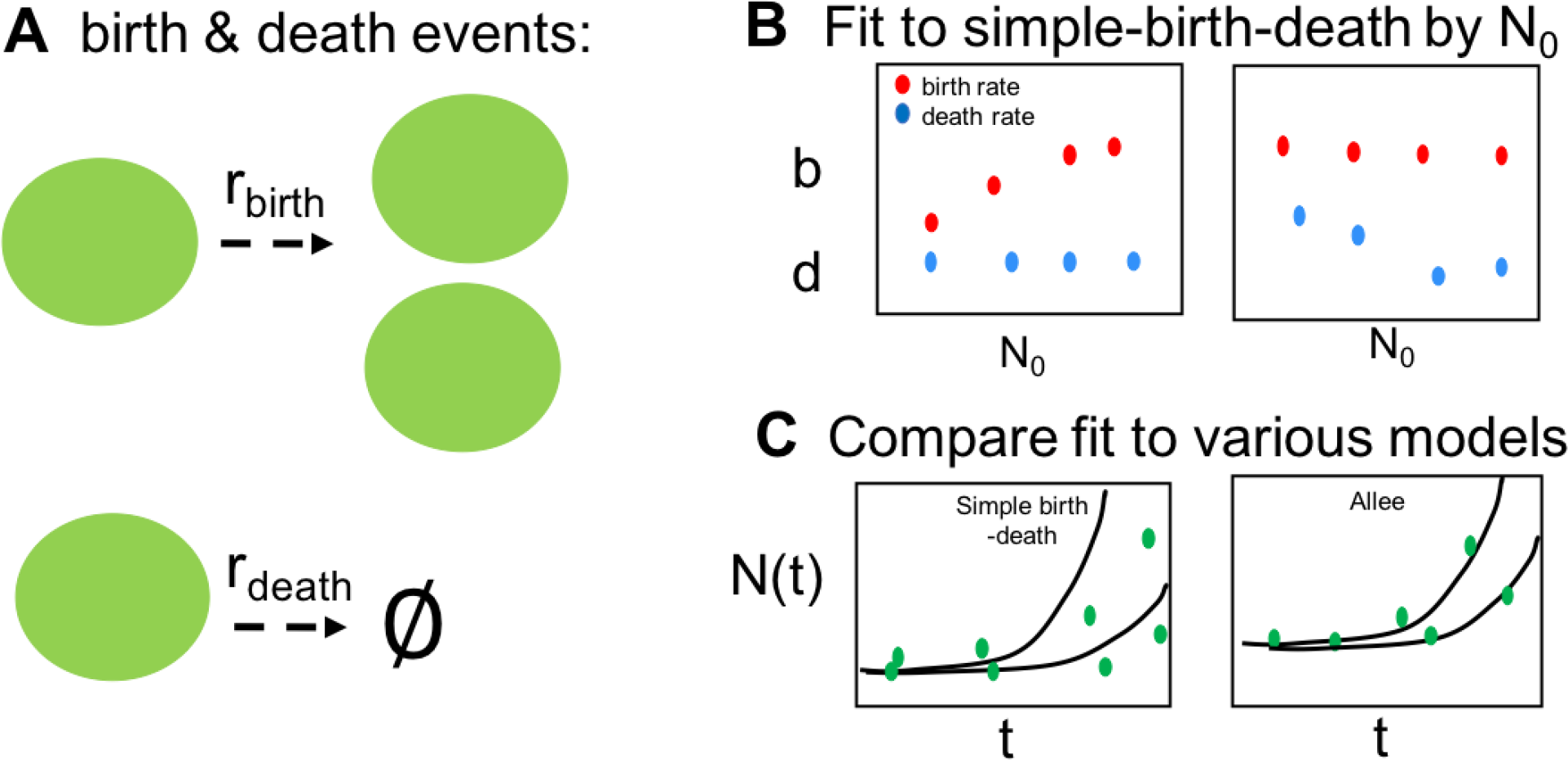
A stochastic model of tumor growth and expected outputs if Allee effect is present. (a) Schematic illustration of generalized stochastic framework where a cell can either give birth or die at a rate given by r_birth_ or r_death_ respectively (b) Schematic of the expected results from fitting of the simplest birth-death model to each data set grouped by initial cell number (N_0_), where, if an Allee effect is present, we expect to observe that either the birth rate constant, b (red), or death rate constant, d (blue), change with initial condition (c) Schematic of the expected outcomes of fitting full data set to the simple birth-death model (left) and a model incorporating an Allee effect (right).

**Figure 3.**
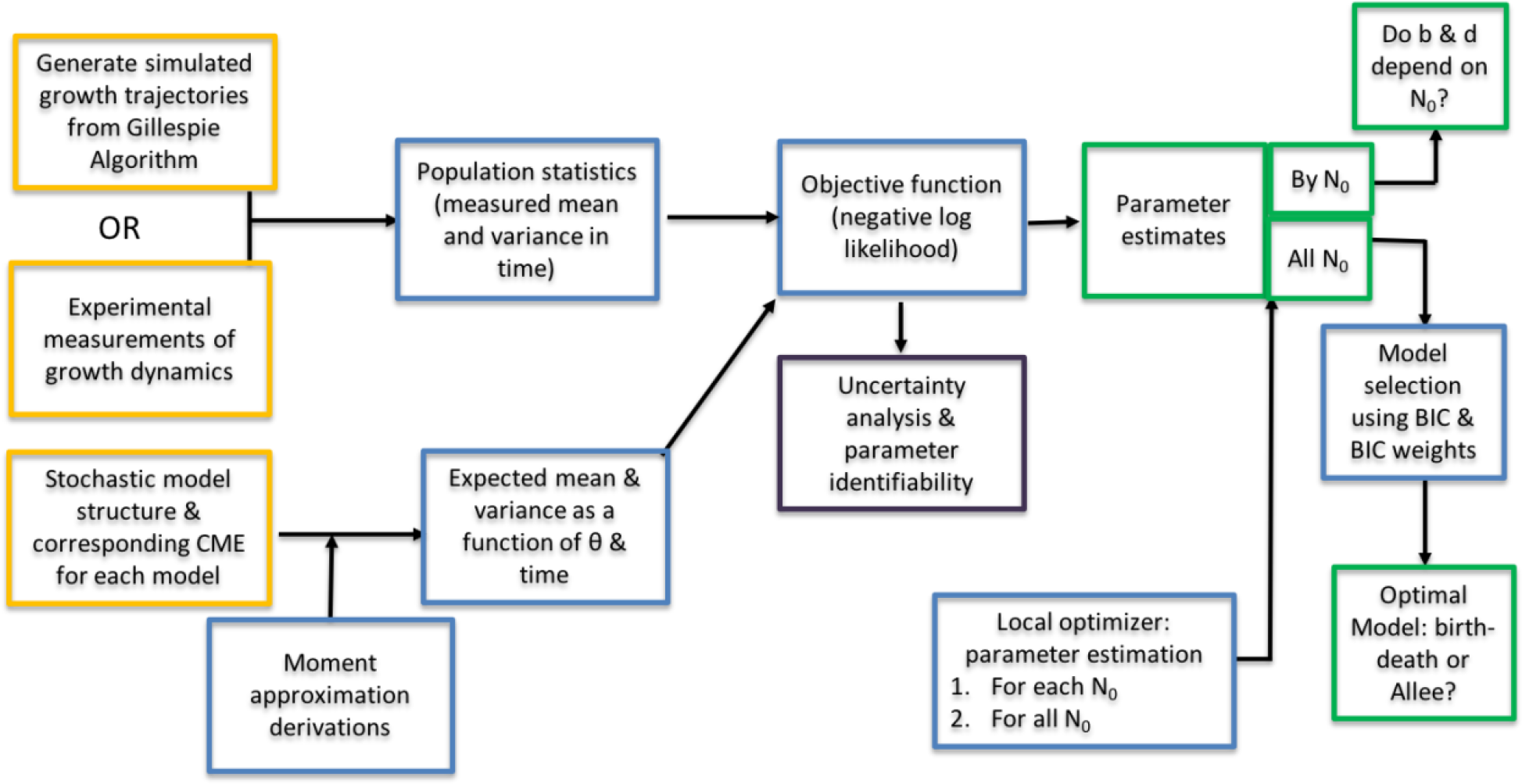
Moment approach for stochastic parameter estimation. Framework for moment approach to derive moments from CME of stochastic process and fit model expected moments to stochastic data.

For each model presented, the generalized framework described above holds and only the birth and death rates (r_birth_ & r_death_) differ for each model based on the hypothesis about the effect of cell number on either birth or death rate. To give an illustrative example of the components of the stochastic model, we explicitly state the reaction rates, the propensities, and the resulting birth and death probabilities for the simple birth-death model. To remain concise, for the remaining six Allee models described we just present the birth and death probabilities for each model.

In the simple birth-death model, the birth rate and death rates are hypothesized to be constant and independent of cell number, n. They are described by rate constants, denoted b & d. The first stochastic model describes a simple birth and death process where the probability of each reaction occurring is given by the rate b or d and the number of cells N.

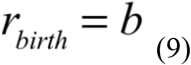

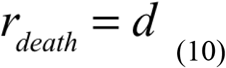

Corresponding to reaction propensities of:

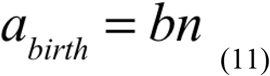

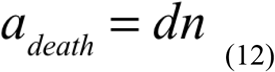

And birth and death probabilities in time- step delta Δt of:

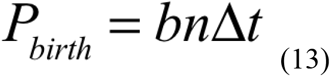

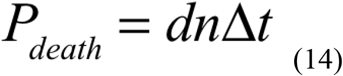

The average behavior in this model corresponds to the exponential growth model (Eq. 1 & 2) where the growth rate constant g is equivalent to the birth rate constant minus the death rate constant (g = b-d). The following set of stochastic models introduce birth and or death rates that are functions of N, corresponding to the hypotheses that the birth and/or death rates are not constant and instead depend on the population size, N. The next family of stochastic models have average behavior that corresponds to the strong Allee effect model in the deterministic case (Eq.4). For the stochastic Allee models, we have to make an assumption of whether the Allee effect acts on the birth probability, the death probability, or on both. (Note that in the deterministic model, these alternatives are indistinguishable, as birth and death rates are combined in an effective growth rate). The strong Allee on birth model assumes that a density-dependent Allee effect lowers the birth probability at N near the Allee threshold, A, and is characterized by the following birth and death rates and probabilities:

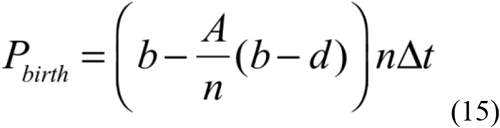

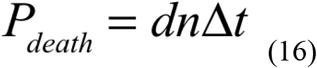

This model hypothesizes that the birth probability is lowered by a factor proportional to the growth rate (b-d), and thus for large n the A/n term in Equation (15) is negligible, but at small n the birth probability is significantly decreased by the Allee term, resulting in a lower birth probability and observed slower net growth.

Alternatively, we could hypothesize that the Allee effect acts to increases the death probability at n near A, resulting in the strong Allee on death model probabilities of:

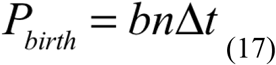

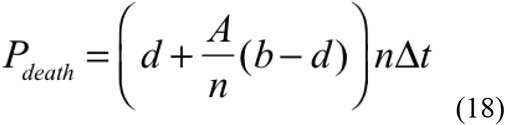

And lastly, we present a model that assumes that the Allee effect term acts equally on both decreasing the birth probability and increasing the death probability for n near A, resulting in:

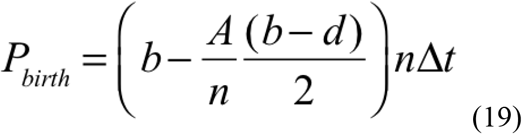

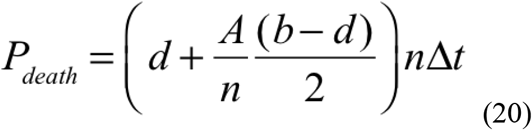

For simplicity, this model assumes that the Allee term acts equally, with half of its effect decreasing the birth rate and half increasing the death rate at n near A. Of course, there could be an infinite number of ways of distributing the Allee threshold onto the birth and death probabilities, and this could have been introduced with an additional fractional parameter. However, for simplicity, we only consider equal partitioning of the Allee effect on both birth and death equally.

The last family of stochastic models corresponds to the extended Allee model (Eq. 5). Again, this model introduces birth and death rate dependencies on n. By the same arguments described for the strong Allee effect model, the extended Allee effect model can manifest itself either on the birth probability only, the death probability only, or the birth and death probabilities equally, leading to the following birth and death probabilities.

If the Allee effect acts on birth only:

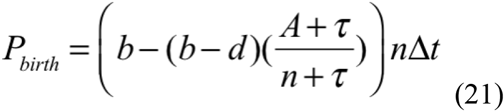

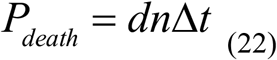

If the Allee effect acts on death only:

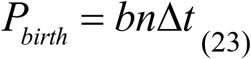

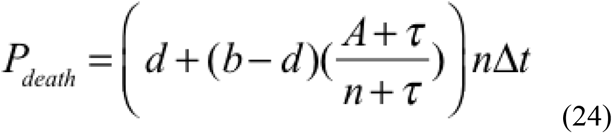

If the Allee effect term acts on birth and death equally:

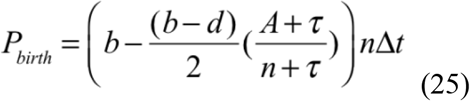

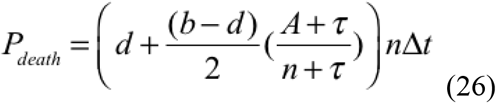

A complete description of each of the above seven stochastic models grouped by the corresponding deterministic model and their assumptions of birth or death mechanism is displayed in Table 1.

**Table 1:** Stochastic growth model families whose average behavior correspond to one of the deterministic growth models. For the Allee model families, within each family the Allee effect can alter birth, death, or both probabilities, leading to distinct mechanistic hypotheses.

| Exponential model family | Strong Allee model family | Extended Allee model family |
| --- | --- | --- |
| **Mean cell number change expressed as deterministic ODE of each family of stochastic models** |  |  |
| $\frac{\partial N}{\partial t} = gN$ | $\frac{\partial N}{\partial t} = gN(1 - \frac{A}{N})$ | $\frac{\partial N}{\partial t} = gN(1 - \frac{A+\tau}{N+\tau})$ |
| **Probabilities of birth & death events to describe stochastic models within each family** |  |  |
| $P(birth) = \Delta t(bN)$<br>$P(death) = \Delta t(dN)$ | Allee effect on birth rate:<br>$P(birth) = \Delta t(bN - (b-d)A)$<br>$P(death) = \Delta t(dN)$ | Allee effect on birth rate<br>$P(birth) = \Delta t(bN - (b-d)N(\frac{A+\tau}{N+\tau}))$<br>$P(death) = \Delta t(dN)$ |
| | Allee effect on death rate:<br>$P(birth) = \Delta t(bN)$<br>$P(death) = \Delta t(dN + (b-d)A)$ | Allee effect on death rate<br>$P(birth) = \Delta t(bN)$<br>$P(death) = \Delta t(dN + (b-d)N(\frac{A+\tau}{N+\tau}))$ |
| | Allee effect on both equally<br>$P(birth) = \Delta t(bN - \frac{(b-d)}{2}A)$<br>$P(death) = \Delta t(dN + \frac{(b-d)}{2}A)$ | Allee effect on both equally<br>$P(birth) = \Delta t(bN - \frac{N(b-d)}{2}(\frac{A+\tau}{N+\tau}))$<br>$P(death) = \Delta t(dN + \frac{N(b-d)}{2}(\frac{A+\tau}{N+\tau}))$ |

To simulate growth trajectories of the stochastic models, we use the Gillespie algorithm (Supp Info S1)(39,40). The above models are used to test the relevance of the Allee effect in cancer cell population growth. The conventional exponential growth model (Eq. 1 & 2) assumes that growth rate (birth rate minus death rate) is constant and independent of initial condition. To test the validity of this assumption in an exploratory analysis, we first fitted each group of trajectories individually for each initial cell number, N_0_. If an Allee effect is present in the data, a systematic increase in the fitted birth rate, b, or decrease in death rate, d, with increasing initial cell number is expected (Figure 2B). We next investigated the relevance of the seven stochastic models by fitting the simulated cell number trajectories from all initial conditions to each stochastic model described above (Equations 9-26) to determine which model structure best describes the observed growth dynamics (Figure 2C).

### Parameter estimation & model selection framework

The parameters of stochastic processes are often inferred using approximate Bayesian computation (41), which require exhaustive stochastic simulations in order to minimize the differences between simulation and experimental data for each parameter set searched. These algorithms require a high number of simulating runs, making them computationally expensive, and instantiating issues of con-convergence and model selection (42). To render inference on the stochastic process (N(t)) feasible, we apply the moment closure approximation method described in Frohlich et al (38) to fit the seven proposed stochastic growth models to experimentally measured growth curves.

#### The chemical master equation of a stochastic growth process

The chemical master equation (CME) describes the change in the probability distribution that the system has any given configuration (in this case number of cells, N) as a function of time. From the CME, the time derivative of the moments, or expected values of n, n^2^,.․n^m^ can be derived. In this framework, we developed stochastic models so that the derivative of the first-order moment corresponds to one of the deterministic models presented (Eqs. 2, 4 &5). The CME describes the probability of their being n cells at time t as a sum of probabilities of a birth, death, or neither event occurring given n−1, n+1, and N cells at time t. An example CME for the birth-death process is:

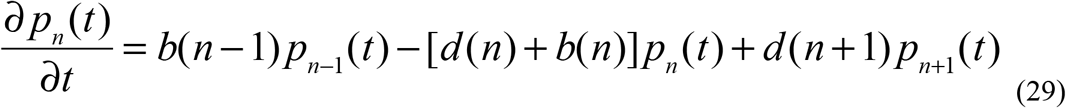

The same procedure was used to derive the CMEs for each of the seven stochastic structural models (Table S2).

#### Derivations of moment-closure approximations from the CME

From longitudinal data of cell number over time (N(t)), with sufficient replicates, we expect to be able to measure the mean and variance in cell number over time. We want to be able to directly compare the mean and variance longitudinal data to the model expected mean and variance in time as a function of the model parameters. We therefore want to derive the first and second moments from the CME. From the CME of each stochastic model (Table S3), the time derivative of the first and second moments were derived according to the procedure outlined in (43)(See Supp. Info S2). Using the definition of variance, the ODEs of the mean and variance for each model can be written in terms of the lower order moments (<n> and <n^2^>, where <.․> denotes the expectation value of the moment)(Table 1). Within each family of models (exponential, strong Allee, and extended Allee, Eqs. 1,2,4, &5) the stochastic forms (Eqs. 9-26) share the same mean ODE corresponding to their deterministic model family, but differ in their variance based on whether the Allee effect is on the birth, death, or both. The time evolution of the variance can be used to properly identify individual rate parameter such as the birth and death rates because the variance in time is proportional not just to the net birth minus death rates, but to the sum of the birth and death rates, as shown in Figure S1.

**Table 2:**
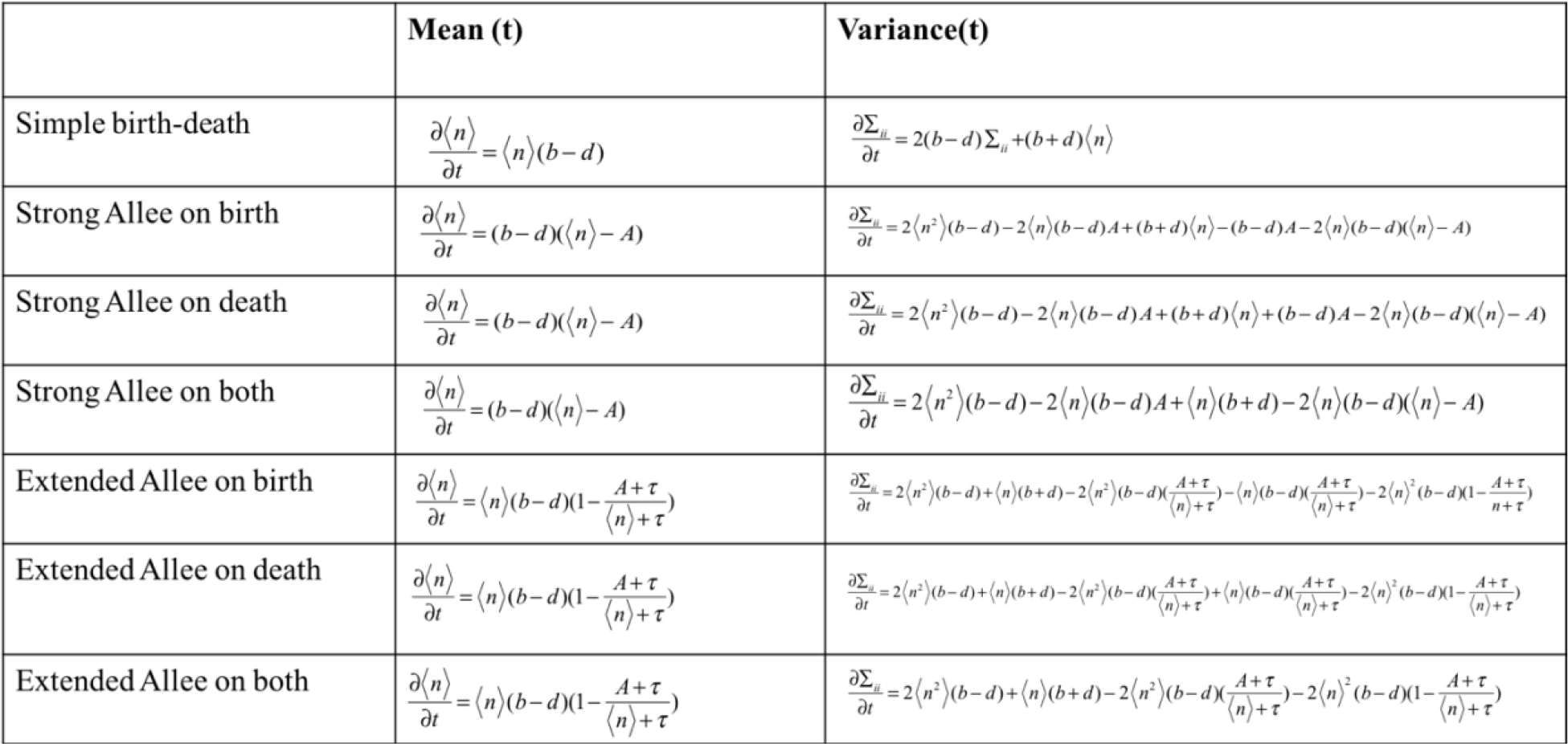
Differential equations of the mean and variance for each stochastic model obtained from the master equation using the moment approach.

|  | Mean (t) | Variance(t) |
| --- | --- | --- |
| Simple birth-death | $\frac{\partial \langle n \rangle}{\partial t} = \langle n \rangle (b - d)$ | $\frac{\partial \Sigma_{ii}}{\partial t} = 2(b - d)\Sigma_{ii} + (b + d)\langle n \rangle$ |
| Strong Allee on birth | $\frac{\partial \langle n \rangle}{\partial t} = (b - d)(\langle n \rangle - A)$ | $\frac{\partial \Sigma_{ii}}{\partial t} = 2\langle n^2 \rangle (b - d) - 2\langle n \rangle (b - d)A + (b + d)\langle n \rangle - (b - d)A - 2\langle n \rangle (b - d)(\langle n \rangle - A)$ |
| Strong Allee on death | $\frac{\partial \langle n \rangle}{\partial t} = (b - d)(\langle n \rangle - A)$ | $\frac{\partial \Sigma_{ii}}{\partial t} = 2\langle n^2 \rangle (b - d) - 2\langle n \rangle (b - d)A + (b + d)\langle n \rangle + (b - d)A - 2\langle n \rangle (b - d)(\langle n \rangle - A)$ |
| Strong Allee on both | $\frac{\partial \langle n \rangle}{\partial t} = (b - d)(\langle n \rangle - A)$ | $\frac{\partial \Sigma_{ii}}{\partial t} = 2\langle n^2 \rangle (b - d) - 2\langle n \rangle (b - d)A + \langle n \rangle (b + d) - 2\langle n \rangle (b - d)(\langle n \rangle - A)$ |
| Extended Allee on birth | $\frac{\partial \langle n \rangle}{\partial t} = \langle n \rangle (b - d) \left(1 - \frac{A + \tau}{\langle n \rangle + \tau}\right)$ | $\frac{\partial \Sigma_{ii}}{\partial t} = 2\langle n^2 \rangle (b - d) + \langle n \rangle (b + d) - 2\langle n^2 \rangle (b - d) \left(\frac{A + \tau}{\langle n \rangle + \tau}\right) - \langle n \rangle (b - d) \left(\frac{A + \tau}{\langle n \rangle + \tau}\right) - 2\langle n \rangle^2 (b - d) \left(1 - \frac{A + \tau}{\langle n \rangle + \tau}\right)$ |
| Extended Allee on death | $\frac{\partial \langle n \rangle}{\partial t} = \langle n \rangle (b - d) \left(1 - \frac{A + \tau}{\langle n \rangle + \tau}\right)$ | $\frac{\partial \Sigma_{ii}}{\partial t} = 2\langle n^2 \rangle (b - d) + \langle n \rangle (b + d) - 2\langle n^2 \rangle (b - d) \left(\frac{A + \tau}{\langle n \rangle + \tau}\right) + \langle n \rangle (b - d) \left(\frac{A + \tau}{\langle n \rangle + \tau}\right) - 2\langle n \rangle^2 (b - d) \left(1 - \frac{A + \tau}{\langle n \rangle + \tau}\right)$ |
| Extended Allee on both | $\frac{\partial \langle n \rangle}{\partial t} = \langle n \rangle (b - d) \left(1 - \frac{A + \tau}{\langle n \rangle + \tau}\right)$ | $\frac{\partial \Sigma_{ii}}{\partial t} = 2\langle n^2 \rangle (b - d) + \langle n \rangle (b + d) - 2\langle n^2 \rangle (b - d) \left(\frac{A + \tau}{\langle n \rangle + \tau}\right) - 2\langle n \rangle^2 (b - d) \left(1 - \frac{A + \tau}{\langle n \rangle + \tau}\right)$ |

For each stochastic model, we confirmed that the mean and variance of a simulated 5000 trajectories with known parameters matched the model mean and variance described in Table 2 (See Supp. Info S3, Fig S2-S8).

#### Maximum likelihood and Bayesian parameter estimation

To infer the parameters of the stochastic models, a maximum likelihood parameter estimation approach was employed using derivations from Frohlich et al (38). The likelihood function assumes that the measured mean and variance of the data at each time point t_k_ is normally distributed around the model predicted first moment (mean cell number (μ(t_k_,θ)) and mean variance in cell number (Σ(t_k_, θ)) with standard deviations for each distribution of mean cell number and variance in cell number given by σ_μ_(θ) and σ_Σ_(θ) respectively. The standard deviation in the first moment, σ_μ_(θ) and the standard deviation in the variance, σ_Σ_(θ), are functions of the parameters θ and were derived by Frohlich et al (38). The likelihood function (Eq. 30) and its corresponding negative log likelihood (Eq.31) are:

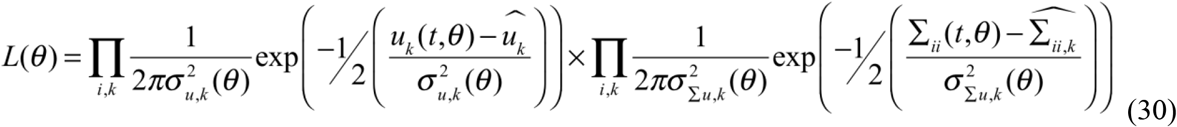

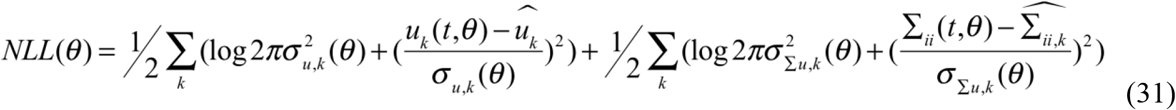

These weigh equally the likelihood of the measured mean and variance of the data from each trajectory over all time points measured. To perform maximum likelihood parameter estimation, we used the fminsearch function in MATLAB to minimize NLL(θ) (Eq. 31). For this optimization, non-negative parameters (rate constants b & d) were log-transformed while parameters able to be negative (extended Allee model Eqs. 21-26, Allee threshold and shape parameter A & τ) were normalized between 0 and 1 over a domain of reasonable values of A and τ. We used the log of the slope of the mean cell number in time as an initial guess for the growth rate (b-d), a death rate of d=0.0005 cells/hour, and an A = 1 or −1 and τ= 2 were used in order to make a conservative initial guess.

#### Uncertainty analysis & parameter identifiability

A key benefit of the moment approach for stochastic parameter estimation is that deriving a system of coupled differential equations enables the use of already established methods for parameter identifiability and uncertainty analysis. To evaluate structural identifiability from each model, a differential algebra approach (44)(45) was used to reveal identifiable combinations of parameters in terms of the output we were able to measure in time, in this case both the mean and the variance of the cell number trajectories in time. (See Supplementary Information S4 for an example of this approach applied to the birth-death model.) This analysis revealed that the parameters in all seven models are uniquely identifiable in theory, i.e. inherent in the model structure.

To ensure that the predicted mean and variance of the models exhibited distinguishable differences from each other, we investigated some illustrative cases of the expected mean and variance for a simple birth-death model, a strong Allee model, and a weak case of the extended Allee model (Figure S9 A & B). Likewise, to ensure the different forms of the stochastic models within each broader class of deterministic models were distinguishable by the expected differences in their variance (Table 1), we display the solutions of the expected mean and variance for strong and weak Allee effects on both birth, death, and equally on both (Figure S10 A-D). This gave us confidence that the candidate models were theoretically distinguishable using the mean and variance of the data collected. To evaluate whether these model parameters were practically identifiable and what the corresponding uncertainty on these model parameters was, the profile likelihood method was used as described in (46). The profile likelihood method evaluates the ability to uniquely identify each parameter individually by ‘profiling’ one parameter at a time, fixing it to a range of values, and fitting for the rest of the parameters. The resulting curvature of likelihood is used to evaluate the uncertainty on the parameter and determine confidence intervals.

### Modeling framework is able to distinguish between different growth models from simulated stochastic trajectories

The parameter estimation and model selection framework were first applied to simulated data based on a stochastic model of intermediate complexity—the strong Allee effect on birth (Eqs. 15 & 16) to demonstrate its capability to expose subtle differences in the underlying growth kinetics of the various models of growth. Using the Gillespie algorithm (39,40) we generated 5000 simulated trajectories from each initial condition of *N*_*0*_= 3, 5, and 10 for the model for strong Allee effect acting on birth.

The stochastic trajectories were sampled every 4 hours corresponding to the time intervals used in the experimental measurements of cell growth, and the mean and variance were calculated at each time point. Artificial noise was added to the calculated mean and variance to test the framework for its ability to distinguish sampling error from stochastic biological variability (Figure 4A & B). The simulated data were fit to the seven candidate models (Eqs. 9-26, Table 2) representing the range of biological hypotheses, with model complexities ranging from 2 to 4 parameters. To identify the most likely underlying model structure from each of the candidate stochastic models, BIC values (47,48) and BIC weights(49) were used for model comparison (See Supp. Info S5). The model selection reveals that the underlying model structure was correctly identified, since the model structure for the strong Allee effect on birth had the lowest BIC value (Figure 4A). The BIC weighting analysis (49) (Supp. Info S5) revealed strong evidence in favor of the strong Allee effect on birth (Figure 4B), indicating the ability of the BIC value to distinguish between overly simple models with 2 parameters and overly complex models with 4 parameters (Figure 4C). In order to ensure the method was not overweighing goodness of fit, the data was down-sampled from the true data collection interval of every 4 hours to every 36 hours to demonstrate that down-sampling changed the magnitudes of the BIC values but did not affect the order of the BIC values of each model relative to one another (Figure S11). The chosen model provided a good fit to the mean and variance in the data (Figure 5 B & C respectively), and the parameter search displayed the expected convergence of accepted parameter values (Figure 5D). Profile likelihoods on parameter distributions demonstrate that each of the parameters are practically identifiable and parameter estimates fall close to the true parameters (Figure 5 E, F, & G). The true parameter values of *b* = 0.00238, d = 0.0005 and *A* = 2 fell within the confidence intervals of the profile likelihood analysis of the fitted parameters of [0.02340, 0.02425] for *b*, [0.00461, 0.00563] for *d* and [1.853, 2.026] for *A*. This confirms that the approach selects appropriate underlying model structure from a set of hypotheses and properly identifies the parameters.

**Figure 4.**
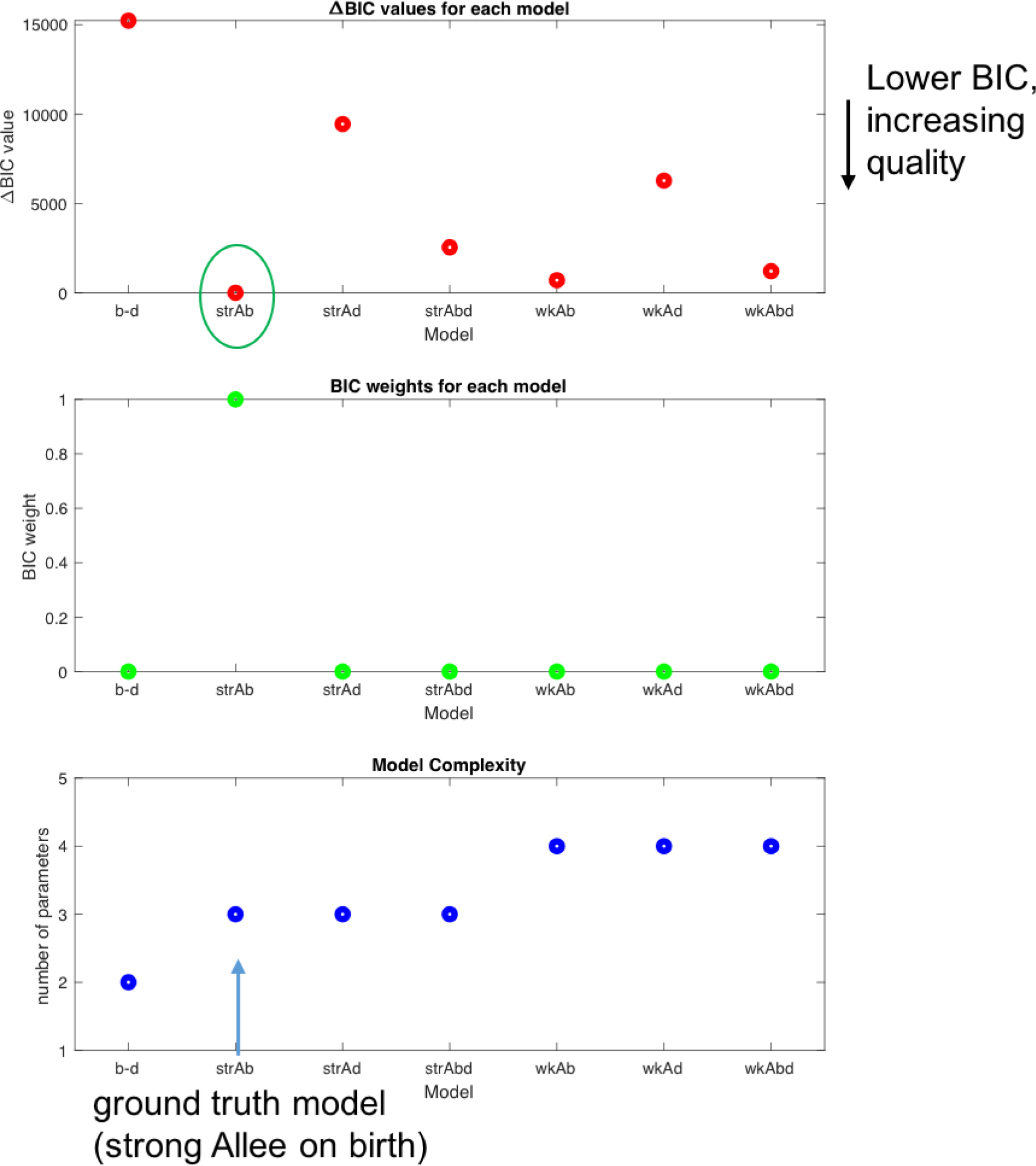
Model selection based on the BIC identifies the “ground truth model” in simulated data. (a) ΔBIC values (BIC_i_-BIC_min_) are plotted for the fit of the simulated data set to each of the seven models [from left-to-right: simple birth-death model (b-d), strong Allee model on birth (strAb), strong Allee model on death (strAd), strong Allee model on birth and death (strAbd), weak extended Allee model on birth (wkAb), weak extended Allee model on death (wkAd), and weak extended Alee model on birth and death (wkAbd)] compared to the minimum BIC value model: the strong Allee model on birth (b) BIC weighting reveals strong evidence to choose the strong Allee model over the other candidate models (c) Number of parameters of each model as a measure of relative complexity of the model.

**Figure 5.**
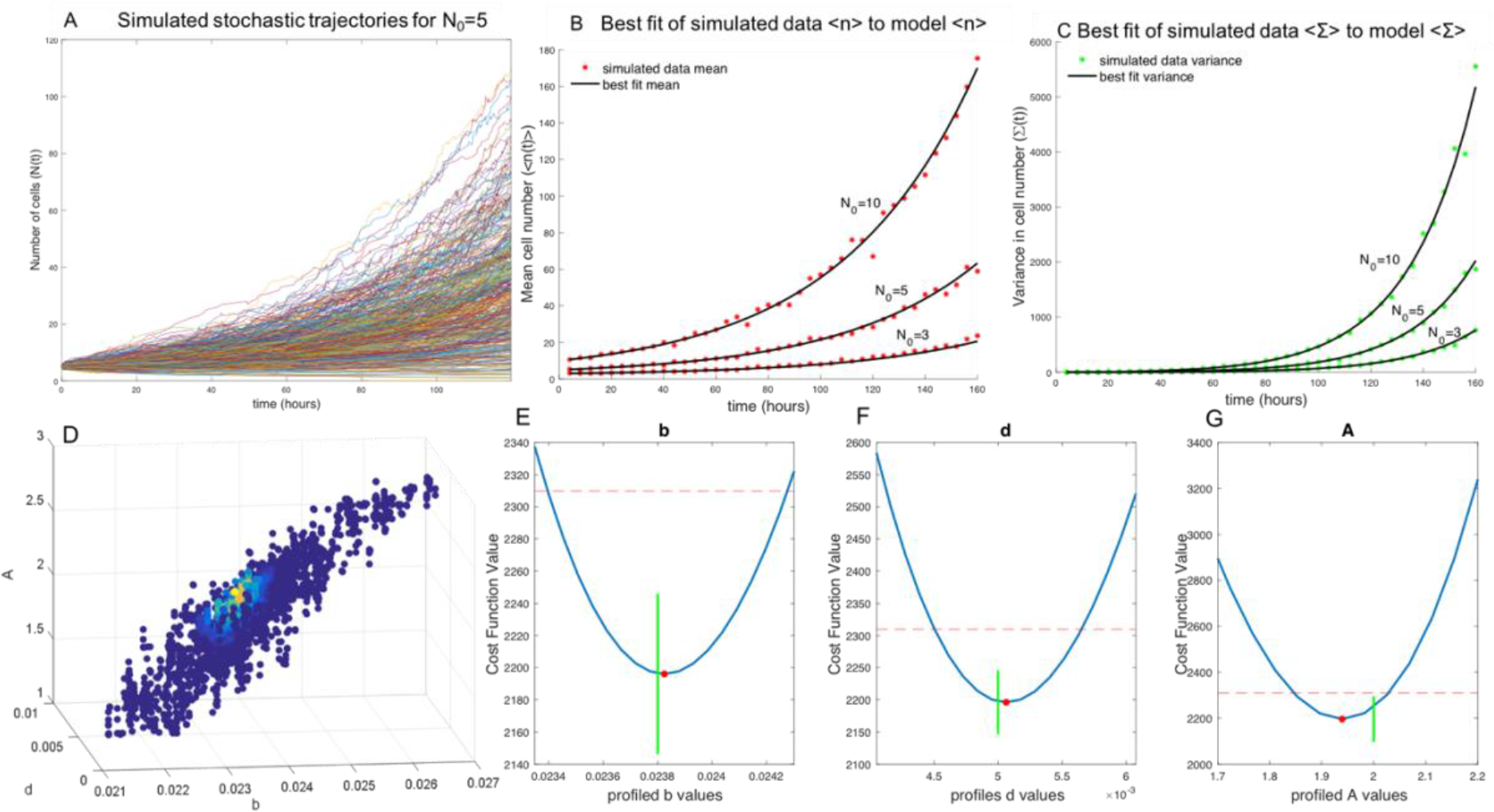
Fit to mean and variance from simulated stochastic data set. (a) Example of stochastic growth model output from 5000 simulated cell number trajectories with initial condition of N_0_=5 and a birth rate of *b* = 0.0238, death rate of *d* = 0.005 and an Allee threshold A =2, revealing the expected variability in growth dynamics apparent at low initial numbers (b) From the simulated stochastic trajectories, we sample time uniformly and measure the mean cell number at each time point for N_0_=3, 5, & 10. (c) The measured variance in cell number at each time point from the uniformly sampled simulated data. (d) Display of parameter space searched, with parameter sets of b, d, and A colored by likelihood, indicating the framework converges on the true parameters (e) Profile likelihood analysis of birth rate parameter estimate (red dot) of *b*=0.0238[0.02340, 0.02425] with true *b* = 0.0238 (green line) (f) Profile likelihood analysis of death rate parameter estimate (red dot) *d* =0.0051 [0.00461, 0.00563] with true *d* = 0.005 (green line) (g) Profile likelihood analysis of Allee threshold parameter estimate (red dot) of *A*=1.9393 [1.853, 2.026] with true *A* = 2(green line).

### Experimental measurement reveals scaling of growth rate with initial cell number

Next we investigated whether the growth of cancer cells *in vitro* is governed by alternative growth models other than the default cell-autonomous proliferation via exponential growth. BT-474 breast cancer cells were seeded at a precise initial cell number ranging from 1 to 20 cells per well of a 96 well plate, and time-lapse microscopy images were collected every 4 hours for replicates wells at each initial condition (20-50 wells per condition) (see *Methods: Cell culture and low cell density seeding*). Example images are shown in Figure 6 A, B, & C. Cell number as function of time was measured for a total of 328 hours (just under 2 weeks) and cell number counts in time were determined using digital image processing for each individual well imaged (See *Methods: Time-lapsed imaging & Image Analysis*).

The true initial cell number N_0_ sorted into each well was confirmed by eye, and wells were binned according to the observed cell number imaged at the first time point. Cell number trajectories of wells with initial cell numbers of 2, 4, and 10 cells are shown in Figure 6D in red, green, and blue respectively. As a preliminary analysis of this data, we fitted each well individually to the exponential growth model (Eqs. 1 & 2) to obtain a distribution of growth rates at each initial condition. The mean growth rates for N_0_ = 2, 4, and 10 respectively were g= 0.00665 +/− 0.00684, 0.00745 +/− 0.00499, and g = 0.00813 +/− 0.00296. Figure 6 displays the average cell number trajectory (Figure 6E) and the normalized growth rate (log(N(t)/N_0_)) (Figure 6F) for the measured data at each time point. These results indicate clear deviations from the simple exponential growth model in which the normalized growth rate (log(N(t)/N_0_)) is expected to be identical for all initial conditions (see Fig. 1C). Instead, growth behavior resembled the characteristic scaling of normalized cell numbers by cell number that is observed for both Allee effect models (Fig. 1F and Fig. 1I). The scaling of average growth rate with initial cell number had been observed by Neufeld et al (3) in their *in vitro* studies of cell culture, providing us with the motivation to further investigate whether an Allee model better describes BT-474 breast cancer cell growth.

To ensure that the observed differences in growth rate at low cell densities were significantly different from what is observed at normal cell culture seeding densities, we sorted N_0_=512 and N_0_=1024 cells and captured 30 growth trajectories from each initial condition. The mean and standard deviation of the growth rates were not significantly different from one another and also significantly higher than the observed low cell density growth rates, with g = 0.0112 +/− 0.00062 and g = 0.0115 +/− 0.00074 for N_0_ = 512 and N_0_ =1024, respectively (Figure S12). The absence of density-dependent growth rates at these higher initial cell numbers may explain why the Allee effect hasn’t been described using standard cell culture seeding densities.

**Figure 6.**
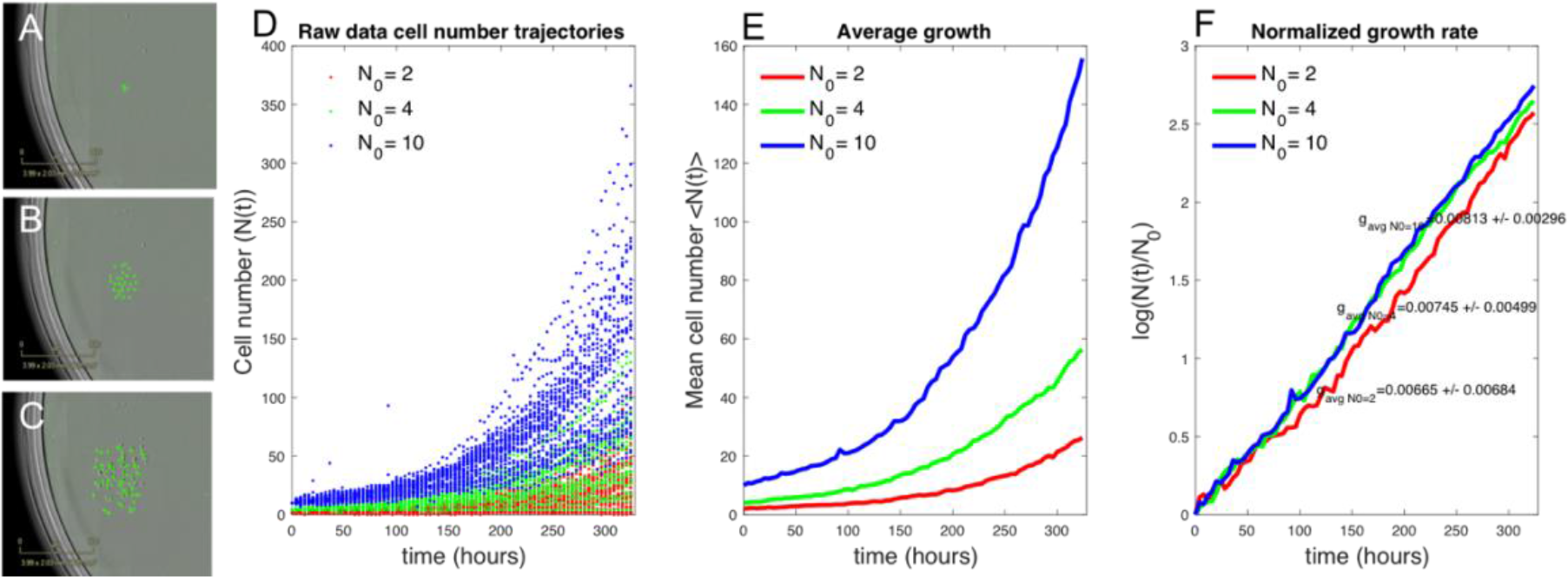
BT-474 cancer cells in culture exhibit growth rate scaling with initial cell density. (a),(b),(c). Representative images from day 1 (a) day 6 (b) and day 14 (c) of BT-474 GFP labeled cells proliferating *in vitro*. (d) Individual cell number trajectories for different N_0_= 2, 4, & 10. (e) Average cell number every 4 hours from each trajectory of N_0_= 2, 4, & 10 (f) Cell number in time normalized by initial cell number in log scale reveals scaling of growth rate by initial cell number, with g = 0.0065 +/−0.00684, 0.00745+/− 0.00499, and 0.00813+/−0.00296 for N_0_= 2, 4, & 10 respectively.

### Fit of experimental data to stochastic growth models reveals Allee effect

The variability in the observed cell number trajectories for a single initial condition is reflected in the experimental measurements of BT-474 cells growing at low initial cell densities (Figure 6D). The variability in cell growth dynamics is expected due to the inherent stochasticity of the birth and death processes which produce a distribution of growth trajectories, as can be seen from the simulated trajectories (Figure 5A). Because stochasticity is more apparent and can be observed experimentally at the low cell numbers (Figure 6D), such dynamics is appropriately modeled by a stochastic rather than a deterministic process. In order to determine whether the preliminary observations of growth rate scaling with the initial cell number could be described by alternative models of cell population growth that consider the Allee effect, the experimental data of BT-474 growth trajectories shown in Figure 6D for initial cell numbers of 2,4, and 10 were calibrated to the seven stochastic models using the stochastic modeling framework presented above (Figure 3).

### Fitting each initial condition separately to the simple birth-death model reveals net growth rate increase with initial cell number

To determine whether birth and/or death rates depend on the initial cell number, we first fit the data for initial cell number of N_0_ = 2, 4, & 10, grouped by initial condition N_0_ individually, to the stochastic simple birth and death model (Eqs. 11-14) using the workflow described in Figure 3.

The results of the fitting to the mean and variance in time to the simple birth-death model for each initial condition are shown in Figure 7 (A-C for the mean and D-F for the variance). Each data set of a single initial condition N_0_ revealed identifiable birth and death rate parameters via profile likelihood analysis (see Figure S13). Birth and death rate maximum likelihood parameter estimates are shown in Figure 7G, with confidence intervals obtained from the profile likelihood analysis (Figure S13). Parameter estimates for birth rates by initial cell number are *b*_*2*_ = 0.00793 [0.00785 0.00794], *b*_*4*_ = 0.00945 [0.0093, 0.0096] & *b*_*10*_ = 0.0113 [0.0112, 0.0114] and for death rates are *d*_*2*_ = 6.67 ×10^−8^ [−0.0005, 0.0001], *d*_*4*_ = 0.0011 [0.0008, 0.0013], & *d*_*10*_ = 0.00286 [0.0025, 0.0028] for N_0_= 2, 4, & 10 respectively. The trend suggests a slight increase in net growth rate (birth rate minus death rate) with initial cell number, as is consistent with the preliminary growth rate analysis by initial cell number (Figure 6F) but inconsistent with the conventional exponential growth hypothesis which should yield the same growth rate (and same birth and death rates), independent of the initial cell number.

**Figure 7.**
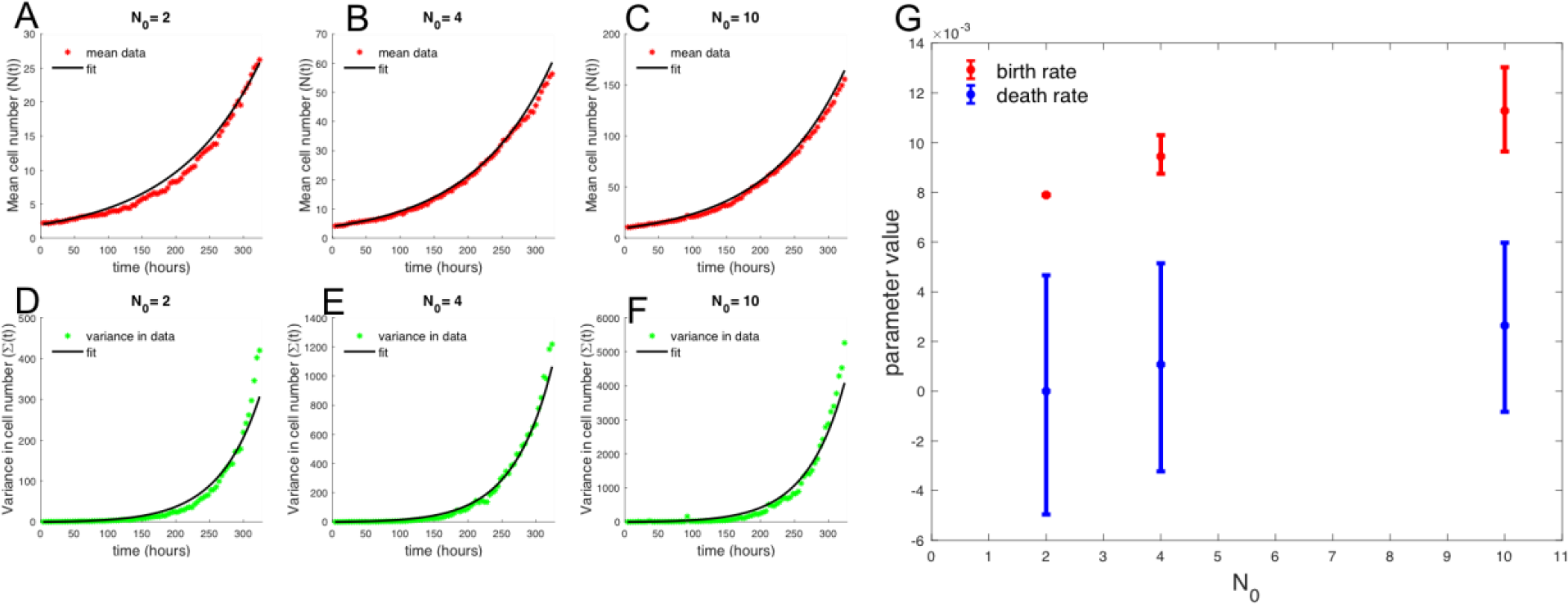
Best fit of each initial cell number means and variance in time to stochastic birth-death model reveals net growth rate increases with initial cell number. (a), (b), (c). Data mean over time compared to best fit to model mean. (d), (e), (f). Data variance in time compared to best fit to model variance. (g) Best fit birth and death rate parameters for the stochastic birth death model fit to each initial condition, with confidence intervals determined from profile likelihoods. Parameter estimates for birth rates by initial cell number are *b*_*2*_ = 0.00793 [0.00785, 0.00794], *b*_*4*_= 0.00945 [0.0093, 0.0096] & *b*_*10*_ = 0.0113 [0.0112, 0.0114] and for death rates are *d*_*2*_ = 6.67 ×10^−18^ [−0.0005, 0.0001], *d*_*4*_ = 0.0011 [0.0008, 0.0013], & *d*_*10*_ = 0.00286 [0.0025, 0.0028] for N_0_= 2, 4, & 10 respectively.

### Fit of low seeding density data to all stochastic models reveals weak Allee effect

The growth data from the initial conditions of *N*_*0*_=2, 4, & 10 were combined and fit to each of the seven candidate models using the moment closure approximation workflow described (Fig 3) (38). The BIC values for each model fit were computed and compared to the minimum BIC value (Figure 8A) and the corresponding BIC weights were calculated (Figure 8B) based on the goodness of fit and the complexity of the model (number of parameters) (Figure 8C). Both the strong and weak Allee effect on birth models have significantly lower BIC values than the null model of the simple birth-death model (Fig 8A), providing strong evidence for the presence of an Allee effect in some form in this data set. Using the BIC weights to evaluate statistical significance between the models revealed that the weak Allee effect on birth is more likely than the strong Allee effect on birth model, with a BIC weight of essentially 1 to 0 for the weak Allee effect on birth versus the strong Allee effect on birth model. The best fit of the weak Allee effect on birth model to the mean and variance of the data is shown in Figure 9 A & B respectively (See Supplementary Figures S14-S19 for the fit of the data to all seven candidate models).

**Figure 8.**
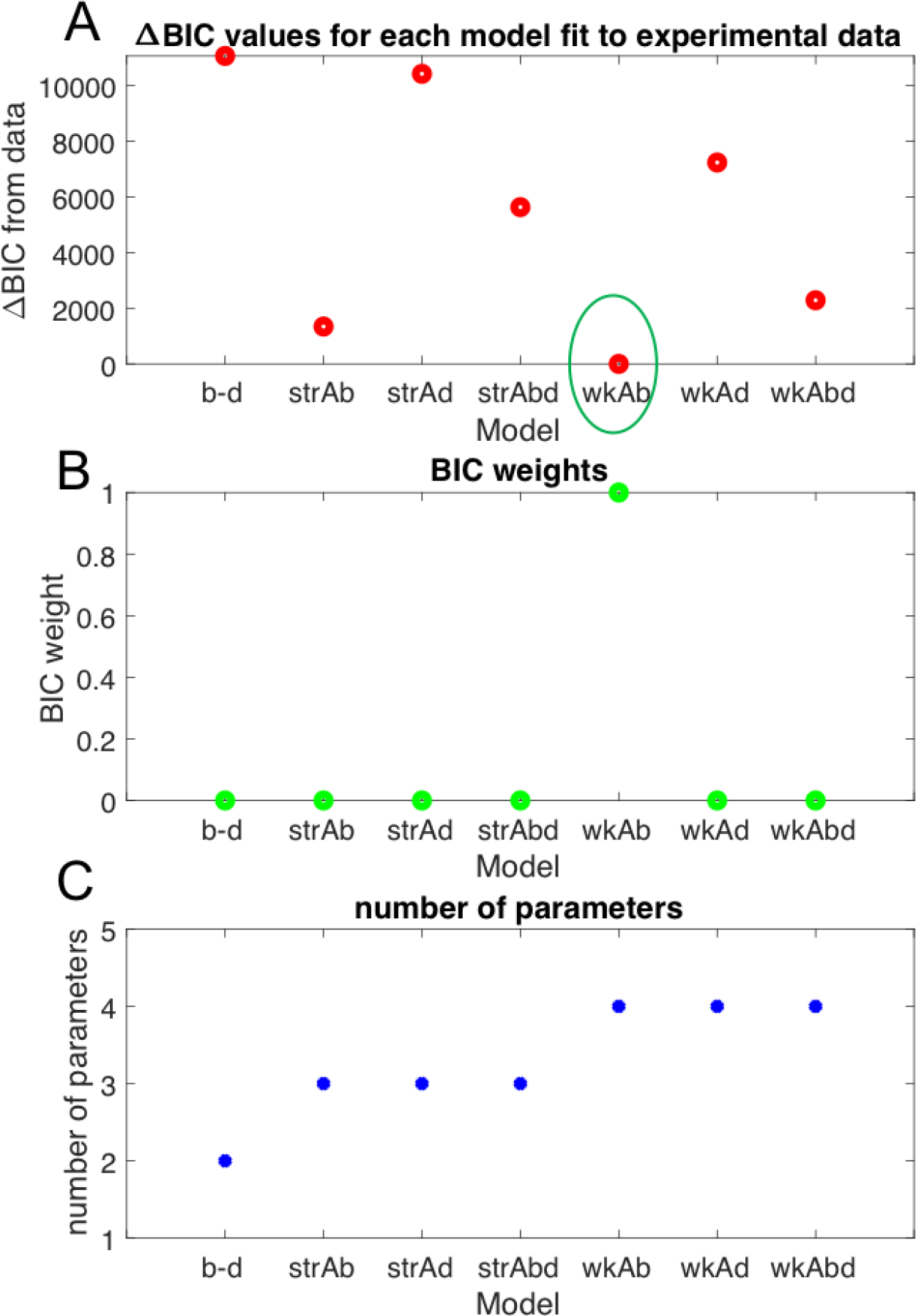
Weak Allee model on birth best describes BT-474 *in vitro* growth data. (a) ΔBIC values for the fit of the data to each of the seven candidate stochastic growth models shows that the weak Allee model on birth exhibits the lowest BIC value. (b) BIC weights for each model indicate that the weak Allee model on birth is significantly better than all other models (c) Number of parameters in each model as a measure of model complexity.

The profile likelihoods used to determine the 95% confidence intervals of the best fitting parameters of *b* = 0.0101[0.010068, 0.010181], *d* = 4.3613 ×10^−5^[−7.27 ×10^−5^, 1.599 ×10^−4^], *A* = −3.1576[-3.8593, −2.4559], and *τ*= 7.480[6.8871, 9.0393] are displayed in Figure 9C, D, E, & F. The discrepancy from the model mean and variance compared to the data is likely because an unbiased approach (as in (38)) was used to fit both the model mean and variance equally, using the likelihood function described in Eq. 30. In theory, the relative weighting of the value of these two outputs could be tuned to reduce the error between the model and measurements in either the mean or variance. The results of model selection for the weak Allee effect model for the BT-474 data indicates that, outside of the effects of demographic stochasticity, any initial cell number is predicted to on average develop into a growing cell population, but the growth rate is expected to be significantly slower at low seeding densities.

**Figure 9.**
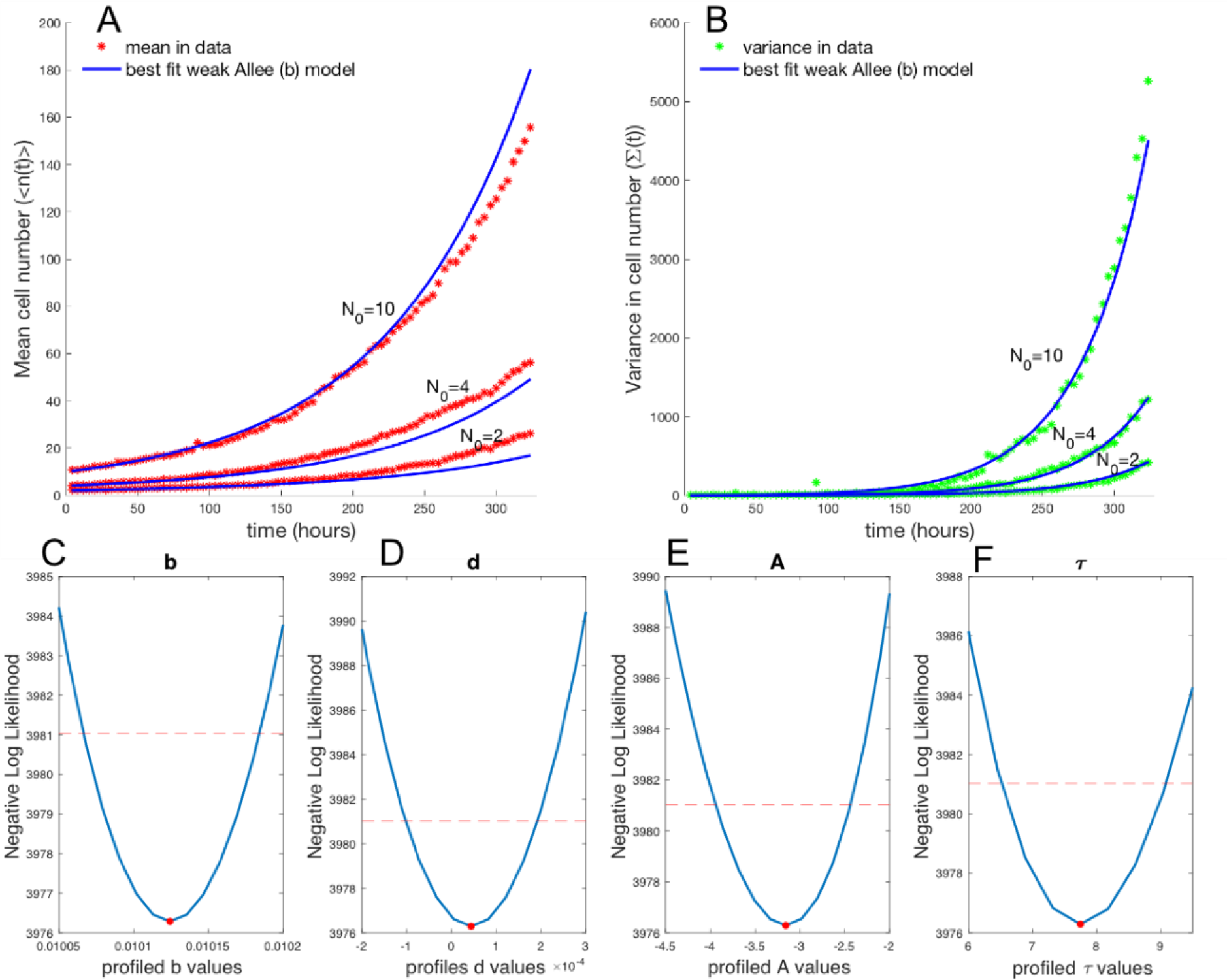
Mean and variance of a subset of data fit to a weak Allee model on birth. (a) Best fit of data mean to model mean displays the model fits the data well over all three initial conditions and over the time course. (b) Best fit of data variance to model variance displays the model fits the data well. (c) Profile likelihood analysis of birth rate around maximum likelihood *b* = 0.0101[0.010068, 0.010181], (d) Profile likelihood analysis of death rate around maximum likelihood *d* =4.3613×10^−5^ [−7.27 ×10^−5^, 1.60 ×10^−4^] (e) Profile likelihood analysis of Allee threshold *A*=−3.1576[−3.8593, −2.4559]. (f) Profile likelihood analysis of the overall shape parameter τ=7.480[6.8871, 9.0393].

## Discussion

The availability of single-cell resolution live imaging of cancer cell growth in a controlled *in vitro* setting starting at the population size of a single cell allowed us to examine in detail the influence of absolute cell number in a cell population on population growth rate (net growth rate = cell birth rate minus death rate) and use mathematical modeling to study the departure from the simple first-order exponential growth kinetics in which the growth rate is proportional to the population size (cell number). Cell-cell interactions, as best known from quorum sensing in bacteria, underlie the cell-number dependence of growth rates. Most work on such dependence have been concerned with the slowing of growth with increasing cell number, e.g. due to approaching the carrying capacity of the cell culture. Here we focus on the initiation of cell growth from a few individual cells and ask whether cooperative behavior, or the Allee effect, as it is known from ecology, can be detected in a departure from exponential growth kinetics as predicted by mathematical models that consider the Allee effect model. Because at the early stages of growth (from one cell or a few) growth kinetics is subjected to stochastic fluctuations due to small cell numbers, we formulated stochastic models that consider the Allee effect. We have demonstrated a framework for testing the relevance of a set of stochastic models of cancer cell growth applied to high-throughput, single cell resolution data.

The seven distinct candidate stochastic models of growth describe various modifications of the exponential growth model by incorporating growth rate dependencies on the size of the population. The average behaviors of these models are examined in the deterministic form, and the corresponding stochastic models that lead to the average behavior are proposed. To test the relevance of the proposed stochastic models, the moment closure approximation method(38) for parameter estimation in stochastic models (Fig. 3) is applied to the high-throughput cell growth data. We first validated our framework by computational simulation of growth trajectories using a model of intermediate complexity. The parameter estimation framework was applied to the simulated data, confirming the ability of the framework to properly identify the underlying model structure and the true parameters. The framework is applied to a data set with a number of replicates from three initial conditions of N_0_ = 2, 4, and 10 BT-474 breast cancer cells. The fit of this growth data reveals that the weak Allee model with the Allee effect decreasing the birth probability at a low cell number, best describes the observed *in vitro* growth dynamics.

The presence of an Allee effect, even in the nutrient and space-rich cell culture setting, implies that cancer cells likely exhibit cooperative growth. The ubiquitous cellular heterogeneity in tumors may suggest that cooperative interactions between distinct subsets of cells must be present in order to maintain the observed heterogeneity(5,50) Recent work by Marusyk et al(6) has found evidence for non-cell autonomous proliferation using a mathematical modeling framework, showing that the null hypothesis of no clonal interactions can be easily rejected in favor of a model that considers a specific clone that helps support the growth of all other clones. Additionally, studies in which clonal diversity has been manipulated by combining clones in culture have demonstrated that the presence of diverse clones is necessary to obtain the observed growth rate achieved in multi-clonal parental cell cultures(51).

Single cell and clonal analysis has enabled the detection of secreted growth inducing factors, such as ILII(6), Wnt1(5), IGFIII(7) and other paracrine factors(8,52) in certain clones which result in an increased growth rate in the surrounding non-producing clones. Bioinformatic analysis of single cell gene expression data has allowed for the identification of specific subsets of cells that produce high levels of certain ligands and coexist in a population with cells that contain high expression levels of the cognate receptors(9,53,54). Prior to single cell analysis capabilities, these types of interactions were not readily detectable from bulk gene expression measurements. In such data the coexpression of a ligand and its cognate receptor in the same sample (a cell population), has by default been interpreted as autocrine signaling(54). Both paracrine and autocrine signaling are likely to play a significant and varying role in tumor growth.

In the field of tumor growth modeling, a few studies have considered the role of the Allee effect and the importance of incorporating it to describe and predict the effects of cooperative growth. Bottger et al(35) developed a stochastic model in which an Allee effect naturally manifests based on assumptions that cancer cells can either exist in a migratory state or a proliferative state. Additional theoretical work has focused on spatial interactions between cancer cells and incorporated the Allee effect in a model for spatial spreading of cancer (34). However, most classical tumor growth models rest on the assumption that early stage growth dynamics match the single exponential growth model (16,23–28). The weak Allee effect revealed in this work provides evidence that descriptions of early stage growth dynamics, which are relevant to progression, relapse, and metastasis, may be improved by taking into account the expected slowing of growth at low cell numbers. Beyond improving predictions of tumor growth and relapse dynamics, a model that considers the Allee effect may help to explain how cancer cell populations are able to go extinct after therapy despite the prediction of the log-kill hypothesis, which states that the probability of a cell being present after treatment, if a tumor is initially large, is greater than zero(55).

While much work in tumor biology has led to an appreciation for cancer as an evolutionary process, a focus on cancer cells as ecosystems of interacting species or subpopulations may yield new insights. The possibility of exploiting ecology for the treatment of tumors based on studies in conservation biology about extinction and control of invasive species has been proposed (13–22), but this is the first work to our knowledge that has explicitly tested for the presence of the Allee effect in a regime in which low cancer cell populations can be measured and fit to a number of stochastic model structures representing different biological hypotheses about the Allee effect. Our finding is consistent with pre-clinical(2) and clinical observations (3,56) of threshold-like behavior of tumor growth or slowed tumor growth following resection. Evidence for the Allee effect is also consistent with evidence of cooperation among cancer cell subclones as has been amply demonstrated (5–7,57–59) An understanding of subpopulation interactions and their molecular mediators that drive the observed Allee effect offer new approaches to manipulate cancer cell growth dynamics in favor of extinction.

This study, which seeks to establish feasibility of detection and mathematical description of the Allee effect by observing growth kinetics, has obvious limitations with respect to biological interpretation of the relevance of results. Most notably we apply the modeling and analysis framework to an *in vitro* data set for a single breast cancer cell line. The *in vitro* system may not faithfully represent *in vivo* growth dynamics, although we expect, and others have shown evidence that (3,56), the Allee effect would only be more pronounced *in vivo*. An *in vitro* setting provides cells with all of the growth factors, nutrients, and space to robustly grow at low cell densities, whereas these factors may be less abundant for tumor cells *in vivo* at a low cell density. While numbers of replicates for each initial condition N_0_ were relatively high (20 to 50 replicates) compared to typical growth studies, an increase in the number of replicates would likely lead to an improvement of the in fit as the variance in the data should be more accurate with increasing sample size. In order to confirm that the Allee effect is a hallmark of tumor growth, a wide range of tumor types will need to be investigated. Additionally, the model presented here is phenomenological; we do not infer the mechanisms by which an Allee effect may be occurring such as in (34,35), nor do we explicitly develop a model of subpopulation interactions as had been done in (6). Future work will focus on investigating the molecular and cellular mechanisms for an Allee effect and developing a model of subpopulation interactions that also considers phenotypic plasticity(60–63).

This work provides a framework for in-depth investigation of mathematical models of stochastic growth that incorporate the Allee effect and shows that an Allee effect model may be more suitable to describing early stage tumor growth dynamics than the exponential model. The potential role of the Allee effect opens a variety of new possibilities for understanding and controlling tumor growth. Biological mechanisms of cooperative growth that may be critical for cell populations to enter a highly proliferative regime need to be further investigated, as these mechanisms may be critical to preventing metastases and tumor relapse.

## Materials & Methods

### Cell culture and low cell density seeding

The human breast cancer cell line BT-474 was used throughout this study. BT-474 is a standard cell line from ATCC. Cell lines were maintained and studied in Dulbecco’s Modified Eagle Medium (DMEM, Thermo Fischer) supplemented with insulin (Gibco) and 10 % fetal bovine serum (Gibco) and 1% Penicillin-Streptomycin (Gibco). A subline of the BT-474 breast cancer cell line was engineered to constitutively express EGFP (enhanced green fluorescent protein) with a nuclear localization signal (NLS). Genomic integration of the EGFP expression cassette was accomplished through the Sleeping Beauty transposon system (64). The EGFP-NLS sequence was obtained as a gBlock from IDT and cloned into the optimized sleeping beauty transfer vector psDBbi-Neo. pSBbie-Neo was a gift from Eric Kowarz (Addgene plasmid #60525)(64). To mediate genomic integration, this two-plasmid system consisting of the transfer vector containing the EGFP-NLS expression cassette and the pCMV(CAT)T7-SB100 plasmid containing the Sleeping Beauty transposase was co-transfected into a BT-474 cell population using Lipofectamine 2000. mCMV(CAT)T7-SB100 was a gift from Zsuzsanna Izsvak (Addgene plasmid # 34879) (65). GFP^+^ cells were collected by fluorescence activated cell sorting. BT-474-EGFPNLS1 cells are maintained in DMEM (Gibco) supplemented with insulin (Sigma Life Science), 10% fetal bovine serum (Fisher) and 200 μg/mL G418 (Caisson Labs). Cells were grown in precoated culture dishes at 37°C in a humidified, 5% CO_2_, 95% air atmosphere. Cells were seeded into the center 60 wells of a 96 well plate (Trueline) at precise initial cell numbers using Fluorescence Activated Cell Sorting (BD Fusion) plate sorting at single cell precision. Plates were kept in the Incucyte Zoom, a combined incubator and time-lapsed microscope. Initial cell seeding numbers were verified by eye at 4x magnification using an image taken within 4 hours from the FACS seeding. Low cell density cultures were allowed to grow in media for 7 days, and were subsequently fed fresh media every 2-3 days for up to two weeks.

### Time lapse imaging

Time-lapse recordings of the cell cultures were performed using the whole-well imaging feature in the Incucyte Zoom. Cells were maintained in the Incucyte at 37°C in humidified 5% CO_2_ atmosphere. Phase contrast and green-channel images were collected every 4 hours for up to 2 weeks.

### Image analysis

Recorded green-channel images were analyzed using the built-in analysis program in the Incucyte Zoom software analysis package. The true initial cell number of each well was confirmed by eye from the images at 4x magnification, and cell number trajectories were binned accordingly. For each 96 well plate, an image processing definition was optimized using the built-in software and confirmed by eye to account for background fluorescence and local bubbles. Wells whose cells died off or did not exhibit any growth and wells with out of focus images were removed from analysis.

## Acknowledgments

This work was supported by the National Institute of Health (R01CA226258, to A.B. and S.H.) and the Texas4000 Foundation Cancer Seed Grant Program (to A.B.). K.J. is grateful for support through an NSF Graduate Research Fellowship. M.K.S. is supported by a postdoctoral fellowship of the German Research Foundation. E. A. B. F. Lima is supported by the National Cancer Institute (NCI) through U01CA174706, and NCI R01CA186193; and the Cancer Prevention Research Institute of Texas (CPRIT) through RR160005.

## References

1. Kobayashi H, Ohkubo M, Narita A, Marasinghe JC, Murao K, Matsumoto T, et al. A method for evaluating the performance of computer- aided detection of pulmonary nodules in lung cancer CT screening: detection limit for nodule size and density. Br J Radiol. 2017;90.

2. Panigrahy D, Edin ML, Lee CR, Huang S, Bielenberg DR, Butterfield CE, et al. Epoxyeicosanoids stimulate multiorgan metastasis and tumor dormancy escape in mice. J Clin Invest. 2012;122(1):178–91.

3. Neufeld Z, Witt W Von, Lakatos D, Wang J, Hegedus B, Czirok A. The role of Allee effect in modelling post resection recurrence of glioblastoma. 2017;1–14.

4. Courchamp F, Berec L, Gascoigne J. Allee Effects in Ecology and Conservation. New York: Oxford University Press; 2008.

5. Cleary AS, Leonard TL, Gestl SA, Gunther EJ. Tumour cell heterogeneity maintained by cooperating subclones in Wnt-driven mammary cancers. Nature. 2014;508(1):113–7.

6. Marusyk A, Tabassum DP, Altrock PM, Almendro V, Michor F. Non-cell-autonomous driving of tumour growth supports sub-clonal heterogeneity. Nature. 2014;514(7520):54–8.

7. Archetti M, Ferraro DA, Christofori G. Heterogeneity for IGF-II production maintained by public goods dynamics in neuroendocrine pancreatic cancer. Proc Natl Acad Sci. 2015;112(6):1833–8.

8. Scheel C, Eaton EN, Li SHJ, Chaffer CL, Reinhardt F, Kah KJ, et al. Paracrine and autocrine signals induce and maintain mesenchymal and stem cell states in the breast. Cell [Internet]. 2011;145(6):926–40. Available from: http://dx.doi.org/10.1016/j.cell.2011.04.029

9. Kumar MP, Du J, Lagoudas G, Jiao Y, Sawyer A, Drummond DC, et al. Analysis of Single-Cell RNA-Seq Identifies Cell-Cell Communication Associated with Tumor Characteristics. Cell Rep. 2018 Nov;25(6):1458–1468.e4.

10. McKenna MT, Weis JA, Brock A, Quaranta V, Yankeelov TE. Precision Medicine with Imprecise Therapy: Computational Modeling for Chemotherapy in Breast Cancer. Transl Oncol [Internet]. 2018;11(3):732–42. Available from: https://doi.org/10.1016/j.tranon.2018.03.009

11. Yankeelov TE, An G, Saut O, Luebeck EG, Popel AS, Ribba B, et al. Multi-scale Modeling in Clinical Oncology: Opportunities and Barriers to Success. Ann Biomed Eng [Internet]. 2016;44(9):2626–41. Available from: https://www.ncbi.nlm.nih.gov/pmc/articles/PMC4983505/pdf/nihms800966.pdf

12. Cloonan N, Forrest ARR, Kolle G, Gardiner BBA, Faulkner GJ, Brown MK, et al. Stem cell transcriptome profiling via massive-scale mRNA sequencing. Nat Methods. 2008;5(7):613–9.

13. Gatenby RA. Population Ecology Issues in Tumor Growth. Cancer Res. 1991;2:2542–8.

14. Basanta D, Anderson ARA, Basanta D, Anderson ARA. Exploiting ecological principles to better understand cancer progression and treatment. Interface Focus. 2013;

15. Korolev KS, Xavier JB, Gore J. Turning ecology and evolution against cancer. Nat Publ Gr [Internet]. 2014;14(5):371–80. Available from: http://dx.doi.org/10.1038/nrc3712

16. West J, Newton PK. Cellular interactions constrain tumor growth. Proc Natl Acad Sci. 2018;

17. Chen K, Pienta KJ. Modeling invasion of metastasizing cancer cells to bone marrow utilizing ecological principles. Theor Biol Med Model. 2011;8(36):1–11.

18. Amend SR, Roy S, Brown JS, Pienta KJ. Ecological paradigms to understand the dynamics of metastasis. Cancer Lett [Internet]. 2016;380(1):237–42. Available from: http://dx.doi.org/10.1016/j.canlet.2015.10.005

19. Amend SR, Pienta KJ. Ecology meets cancer biology: The cancer swamp promotes the lethal cancer phenotype. Oncotarget. 2015;6(12).

20. Axelrod R, Pienta KJ. Cancer as a Social Dysfunction — Why Cancer Research Needs New Thinking. Mol Cancer Res. 2018;16(September):2018–20.

21. Mcgregor N, Axelrod R, Axelrod DE. Ecological Therapy for Cancer: Defining Tumors Using an Ecosystem Paradigm Suggests New Opportunities for Novel Cancer Treatments. Transl Oncol. 2008;1(4):158–64.

22. Han J, Jun Y, Hyun S, Hoang H, Jung Y, Kim S, et al. Rapid emergence and mechanisms of resistance by U87 glioblastoma cells to doxorubicin in an in vitro tumor microfluidic ecology. Proc Natl Acad Sci. 2016;113(50):14283–8.

23. Pacheco E. A review of models for cancer chemotherapy based on Optimal Control. INESC-ID Tech Rep. 2016;1–30.

24. Benzekry S, Lamont C, Beheshti A, Tracz A, Ebos JML, Hlatky L, et al. Classical Mathematical Models for Description and Prediction of Experimental Tumor Growth. PLoS Comput Biol. 2014;10(8).

25. Lima EABF, Oden JT, Hormuth D, Yankeelov TE, Almeida RC. Selection, calibration, and validation of models of tumor growth. Math Model Appl Sci. 2016;

26. Norton L. A Gompertzian Model of Human Breast Cancer Growth. Cancer Res. 1988;7067–71.

27. Speer JF, Petrosky VE, Retsky MW, Wardwell RH. A Stochastic Numerical Model of Breast Cancer Growth That Simulates Clinical Data. Cancer Res. 1984;(44):4124–30.

28. Winsor CP. The Gompertz curve as a growth curve. Proc Natl Acad Sci. 1932;18(1).

29. Bose I, Prafulla A, Road C, Pal M, Prafulla A, Road C, et al. Allee dynamics: Growth, extinction and range expansion. arXiv. 2017;(1):1–9.

30. Vieira R, Ribeiro FL, Souto A. Models for Allee effect based on physical principles. J Theor Biol [Internet]. 2015;385:143–52. Available from: http://dx.doi.org/10.1016/j.jtbi.2015.08.018

31. Duncan RP, Blackburn TM, Rossinelli S, Bacher S. Quantifying invasion risk: The relationship between establishment probability and founding population size. Methods Ecol Evol. 2014;5(11):1255–63.

32. Wittmann M, Gabriel W, Metzler D. Genetic Diversity in Introduced Populations with an Allee Effect. Genet Soc Am. 2014;198(September):299–310.

33. Rodriguez-brenes IA, Komarova NL, Wodarz D. Tumor growth dynamics: insights into evolutionary processes. Trends Ecol Evol [Internet]. 2013;28(10):597–604. Available from: http://dx.doi.org/10.1016/j.tree.2013.05.020

34. Sewalt L, Harley K, Heijster P Van, Balasuriya S. Influences of Allee effects in the spreading of malignant tumours. J Theor Biol [Internet]. 2016;394:77–92. Available from: http://dx.doi.org/10.1016/j.jtbi.2015.12.024

35. Böttger K, Hatzikirou H, Voss-böhme A. An Emerging Allee Effect Is Critical for Tumor Initiation and Persistence. PLoS Comput Biol. 2015;1–14.

36. Greene JM, Levy D, Herrada SP, Gottesman MM. Mathematical Modeling Reveals That Changes to Local Cell Density Dynamically Modulate Baseline Variations in Cell Growth and Drug Response. Cancer Res. 2016;76(10):2882–91.

37. Konstorum A, Hillen T, Lowengrub J. Effect, Feedback Regulation in a Cancer Stem Cell Model can Cause an Allee Affect. Bull Math Biol. 2016;78(4):754–85.

38. Fröhlich F, Thomas P, Kazeroonian A, Theis FJ. Inference for Stochastic Chemical Kinetics Using Moment Equations and System Size Expansion. PLoS Comput Biol. 2016;12(7):1–28.

39. Gillespie DT. The chemical Langevin equation. J Chem Phys. 2014;297(2000).

40. Gillespie DT. Exact Stochastic Simulation of Coupled Chemical Reactions. J Phys Chem. 1977;81(25):2340–61.

41. Beaumont MA, Zhang W, Balding DJ. Approximate Bayesian Computation in Population Genetics. Genet Soc Am. 2002;162(December):2025–35.

42. Robert CP, Cornuet J, Marin J, Pillai NS. Lack of confidence in approximate Bayesian computation model choice. Proc Natl Acad Sci. 2011;108(37).

43. Houchmandzadeh B. Extracting moments from Master Equations. ArXiv. 2009;1(2):1–14.

44. Meshkat N, Sullivant S, Eisenberg M. Identifiability Results for Several Classes of Linear Compartment Models. Bull Math Biol. 2015;77(8):1620–51.

45. Brouwer AF, Meza R, Eisenberg MC, Arbor A. A systematic approach to determining the identifiability of multistage carcinogenesis models. Risk Anal. 2018;37(7):1375–87.

46. Raue A, Kreutz C, Maiwald T, Bachmann J, Schilling M, Klingmüller U, et al. Structural and practical identifiability analysis of partially observed dynamical models by exploiting the profile likelihood. Bioinformatics. 2009;25(15):1923–9.

47. Raftery A. Bayes Factors and BIC. Sociol Methods Res. 1999;27(3):411–27.

48. Loos C, Moeller K, Fröhlich F, Hucho T, Hasenauer J. A Hierarchical, Data-Driven Approach to Modeling Single-Cell Populations Predicts Latent Causes of Cell-To-Cell Variability. Cell Syst. 2018;6(5):593–603.e13.

49. Wagenmakers E, Farrell S. AIC model selection using Akaike weights. Psychon Bull Rev. 2004;11(1):192–6.

50. Brock A, Chang H, Huang S. Non-genetic heterogeneity — a mutation-independent driving force for the somatic evolution of tumours. Nat Rev Genet [Internet]. 2009;10(5):336–42. Available from: http://www.nature.com/doifinder/10.1038/nrg2556

51. Wangsa D, Braun R, Schiefer M, Gertz EM, Bronder D, Padilla-nash IQHM, et al. The evolution of single cell-derived colorectal cancer cell lines is dominated by the continued selection of tumor-specific genomic imbalances, despite random chromosomal instability. Carcinogenesis. 2018;(June):1–13.

52. Hoelzinger DB, Demuth T, Berens ME. Autocrine factors that sustain glioma invasion and paracrine biology in the brain microenvironment. J Natl Cancer Inst. 2007;99(21):1583–93.

53. Zhou JX, Taramelli R, Pedrini E, Knijnenburg T, Huang S. Extracting Intercellular Signaling Network of Cancer Tissues using Ligand-Receptor Expression Patterns from Whole-tumor and Single-cell Transcriptomes. 2017;(July):1–15.

54. Graeber TG, Eisenberg D. Bioinformatic identification of potential autocrine signaling loops in cancers from gene expression profiles. Nat Genet. 2001;29(3):295–300.

55. Poleszczuk J, Enderling H. The Optimal Radiation Dose to Induce Robust Systemic Anti-Tumor Immunity. Int J Mol Sci. 2018;19(11).

56. Spiteri I, Caravagna G, Cresswell GD, Vatsiou A, Nichol D, Acar A, et al. Evolutionary dynamics of residual disease in human glioblastoma. Oxford Univ Press. 2018;

57. Axelrod R, Axelrod DE, Pienta KJ. Evolution of cooperation among tumor cells. Proc Natl Acad Sci. 2006;103(36):13474–9.

58. An MW, Dong X, Meyers J, Han Y, Grothey A, Bogaerts J, et al. Evaluating continuous tumor measurement-based metrics as phase II endpoints for predicting overall survival. J Natl Cancer Inst. 2015;107(11):1–7.

59. Brown JL, Russelll PJ, Philips J, Wotherspoon J, Raghavan D, Prince R, et al. Clonal analysis of a bladder cancer cell line: tumour heterogeneity experimental model of. Br J Cancer. 1990;61:369–76.

60. Pisco AO, Huang S. Non-genetic cancer cell plasticity and therapy-induced stemness in tumour relapse: ‘What does not kill me strengthens me.’ Br J Cancer [Internet]. 2015;112(11):1725–32. Available from: http://dx.doi.org/10.1038/bjc.2015.146

61. Zhou JX, Pisco AO, Qian H, Huang S. Nonequilibrium population dynamics of phenotype conversion of cancer cells. PLoS One. 2014;9(12):1–19.

62. Gupta PB, Fillmore CM, Jiang G, Shapira SD, Tao K, Kuperwasser C, et al. Stochastic state transitions give rise to phenotypic equilibrium in populations of cancer cells. Cell [Internet]. 2011;146(4):633–44. Available from: http://dx.doi.org/10.1016/j.cell.2011.07.026

63. Jolly MK, Tripathi SC, Somarelli JA, Hanash SM, Levine H. Epithelial/mesenchymal plasticity: how have quantitative mathematical models helped improve our understanding? Mol Oncol. 2017;11(7):739–54.

64. Kowarz E, Loescher D, Marschalek R. Optimized Sleeping Beauty transposons enable robust stable transgenic cell lines. Biotechnol J. 2015;41:647–53.

65. Mátés L, Chuah MKL, Belay E, Jerchow B, Manoj N, Acosta-Sanchez A, et al. Molecular evolution of a novel hyperactive Sleeping Beauty transposase enables robust stable gene transfer in vertebrates. Nat Genet. 2009;41(6):753–61.

