## Supplementary Information for "Cancer cell population growth kinetics at low densities deviate from the exponential growth model and suggest an Allee effect" (from another version of this preprint: version 1, 2019-03-21)

### Supporting Information

| Hypothesis | Model | Parameters | Behavior |
|---|---|---|---|
| No Allee effect | $\frac{\partial N}{\partial t} = gN$ | g = intrinsic growth rate | Cells exhibit cell autonomous proliferation and will grow at a constant rate regardless of cell number |
| Strong Allee effect | $\frac{\partial N}{\partial t} = gN(1 - \frac{A}{N})$ | A = Allee threshold | If the number of cells is below the threshold A, the population will go extinct. If N is near A, the growth rate will be slowed until N>>A when the growth rate will approach g |
| Weak/strong Allee effect | $\frac{\partial N}{\partial t} = gN(1 - \frac{A+\tau}{N+\tau})$ | **τ** = overall shape parameter, as **τ** increases, per capita growth flattens | When A is positive, strong Allee effect, when **τ**>\|A\| and A is negative, the population will grow more slowly when N is near A, but will not go extinct |

Table S1. Deterministic model structures to describe three distinct tumor growth dynamic model hypotheses.

#### S1. Stochastic model simulation using the Gillespie Algorithm

To evolve the stochastic model forward, we implement the Gillespie algorithm (1)(2) by initializing the number of cells to begin with and the rate parameters that describe the probability of either a birth or death event. The Monte Carlo step is performed to generate random numbers that determine the next event (either a birth or a death) to occur as well as the time interval until that event occurs. The probability of a given event to be chosen is proportional to the reaction propensity and the time interval to the next event is exponentially distributed with mean of the reciprocal of the sum of the probability of any of the events occurring. This process is described below:

$$r1 = rand \in [0,1]$$
$$r2 = rand \in [0,1]$$
$$r1 < \frac{P(birth)}{P(birth) + P(death)}$$
$$N = N + 1$$
$$else$$
$$N = N - 1$$
$$\Delta t = \frac{-\log(r2)}{P(birth) + P(death)}$$

Where P(birth) and P(death) are specific to the stochastic model structure (See Table S2 for definitions that correspond to each model). These steps were repeated for up to 5000 repetitions. Because the time step was probabilistic, each stochastic trajectory was sampled to obtain a uniform time interval where the number of cells was recorded at each time based on the number at the event equal to or before the interval.

| Stochastic Model | Chemical Master Equation |
|---|---|
| Birth-death model | $\frac{\partial p_n(t)}{\partial t} = b(n-1)p_{n-1}(t) - [d(n)+b(n)]p_n(t) + d(n+1)p_{n+1}(t)$ |
| Strong Allee on birth model | $\frac{\partial p_n(t)}{\partial t} = [b(n-1)-(b-d)A]p_{n-1}(t) - [d(n)+b(n)-(b-d)A]p_n(t) + d(n+1)p_{n+1}(t)$ |
| Strong Allee on death model | $\frac{\partial p_n(t)}{\partial t} = b(n-1)p_{n-1}(t) - [d(n)+(b-d)A+b(n)]p_n(t) + [d(n+1)+(b-d)A]p_{n+1}(t)$ |
| Strong Allee on birth & death model | $\frac{\partial p_n(t)}{\partial t} = [b(n-1)-\frac{(b-d)}{2}A]p_{n-1}(t) - [d(n)+b(n)]p_n(t) + [d(n+1)+\frac{(b-d)}{2}A]p_{n+1}(t)$ |
| Weak Allee on birth model | $\frac{\partial p_n(t)}{\partial t} = [b(n-1)-(b-d)(n-1)\frac{(A+\tau)}{(n-1+\tau)}]p_{n-1}(t) - [d(n)+b(n)-(b-d)(n)\frac{(A+\tau)}{(n+\tau)}]p_n(t) + d(n+1)p_{n+1}(t)$ |
| Weak Allee on death model | $\frac{\partial p_n(t)}{\partial t} = b(n-1)p_{n-1}(t) - [d(n)+(b-d)\frac{(A+\tau)}{(n+\tau)}+b(n)]p_n(t) + [d(n+1)+(b-d)(n+1)\frac{(A+\tau)}{(n+1+\tau)}]p_{n+1}(t)$ |
| Weak Allee on birth & death model | $\frac{\partial p_n(t)}{\partial t} = [b(n-1)-\frac{(b-d)(n-1)}{2}\frac{(A+\tau)}{(n-1+\tau)}]p_{n-1}(t) - [d(n)+b(n)]p_n(t) + [d(n+1)+\frac{(b-d)(n+1)}{2}\frac{(A+\tau)}{(n+1+\tau)}]p_{n+1}(t)$ |

Table S2. Chemical Master Equations to describe each of the stochastic model structures.

#### S2. Derivation of the moment-closure approximation for the first moment of the birth-death model

Starting from the CME for the birth-death model below, we apply the $\sum n^m$th operator (in the case of the first moment m=1) over the time derivative of the probability of their being n cells at time t to obtain the dime derivative of the expectation of n.

$$\frac{dp_n(t)}{dt} = b(n-1)p_{n-1} - [d(n)+b(n)]p_n + d(n+1)p_{n+1}$$

$$\sum_{n=-\infty}^{n=\infty} np_n(t) = \langle n \rangle$$

$$\frac{d\left(\sum_{n=-\infty}^{n=\infty} np_n(t)\right)}{dt} = \frac{d\langle n \rangle}{dt}$$

We apply the $\sum n^m$th operator to each term on the RHS of the CME. We can then use the fact that the summation from -∞ to ∞ is the same over n-1, n+1, and n to transform the $p_{n+1}$ and $p_{n-1}$ to $p_n$s by transforming each term on the RHS so that n-1= n, and substituting into each multiplicative term accordingly as shown below:

$$\frac{d\left(\sum_{n=-\infty}^{n=\infty} np_n(t)\right)}{dt} = \sum_{n=-\infty}^{n=\infty} nb(n-1)p_{n-1} - \sum_{n=-\infty}^{n=\infty} n[d(n)+b(n)]p_n + \sum_{n=-\infty}^{n=\infty} nd(n+1)p_{n+1}$$

$$\sum_{n=-\infty}^{n=\infty} n = \sum_{n=-\infty}^{n=\infty} n-1$$

$$\frac{d\left(\sum_{n=-\infty}^{n=\infty} np_n(t)\right)}{dt} = \sum_{n=-\infty}^{n=\infty} (n+1)b(n)p_n - \sum_{n=-\infty}^{n=\infty} n[d(n)+b(n)]p_n + \sum_{n=-\infty}^{n=\infty} (n-1)d(n)p_n$$

Factoring out pn and applying like terms:

$$\frac{d\left(\sum_{n=-\infty}^{n=\infty} np_n(t)\right)}{dt} = \sum_{n=-\infty}^{n=\infty} b(n)p_n - \sum_{n=-\infty}^{n=\infty} d(n)p_n$$

Applying the $\sum n^m$th operator gives the time derivative of the expected first moment in terms of the first moment of n itself. In this case, we can solve this analytically to obtain the moment-approach approximation for the mean cell number of the stochastic birth-death process described by exponential growth with a growth rate equal to the birth rate minus the death rate.

$$\frac{d\langle n\rangle}{dt} = (b-d)\langle n\rangle$$

$$\langle n\rangle(t) = \langle n\rangle_0 e^{(b-d)t}$$

This same derivation(3) was repeated to find up to the 4th moment, and the definition of the variance was used to derive the expected time-derivative of the variance use to identify the magnitude of the birth and death rate.

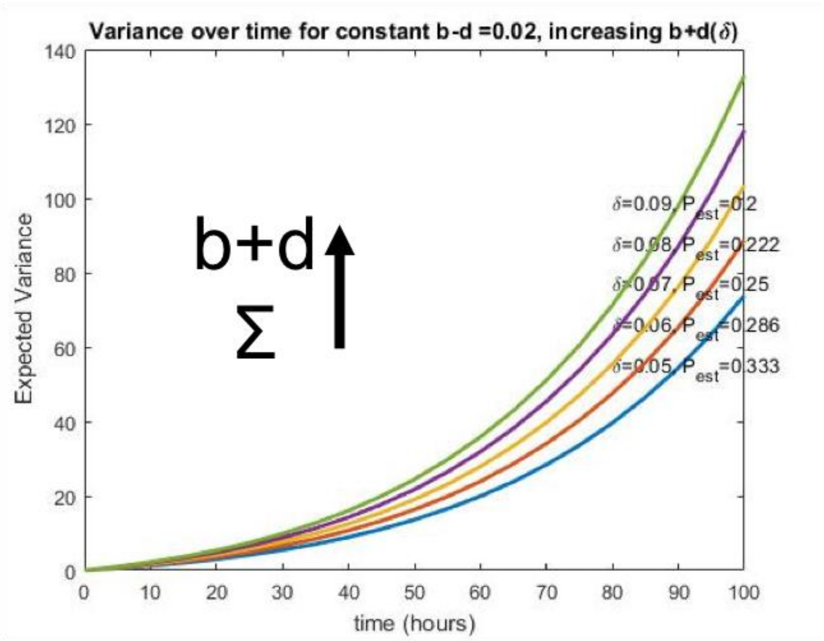


**Figure S1. Illustrative example demonstrates that increasing magnitude of birth and death rate parameters increases variance in time, enabling the identifiability of b+d.** The net growth (b-d) was held constant and the magnitude of b and d were simultaneously increased in order to demonstrate the effect on the time evolution of the variance. This example is used to explain intuitively how the measurement of variance in time enables the proper identification of the b and d parameters uniquely, even while the time evolution of the mean cell number remains constant.

#### S3. Confirmation that derivations of mean and variance for each model match the mean and variance from simulated data with known parameters

The moment-approximation derivations from the CME were confirmed to match the measured moments from simulated data from the Gillespie algorithm. In Figure S2, five thousand trajectories are simulated from the stochastic birth-death model (Eq. 5 &6) with an initial cell number on $N_0$=5 and a birth rate of b = 0.0238 cells/ hour and a death rate of d = 0.005 cells/hour (Figure 4A). The stochastic simulation trajectories were sampled every 4 hours, and the mean and variance in cell number were calculated at each time point (Figure 4B). Figure 4C shows the measured mean from simulated data and the expected mean are a near perfect match, and Figure 4D shows the measured and expected variance are as well. See Supplementary Figures S2-7 for confirmation that the expected mean and variance from the remaining six stochastic models match the measured mean and variance from simulated data.

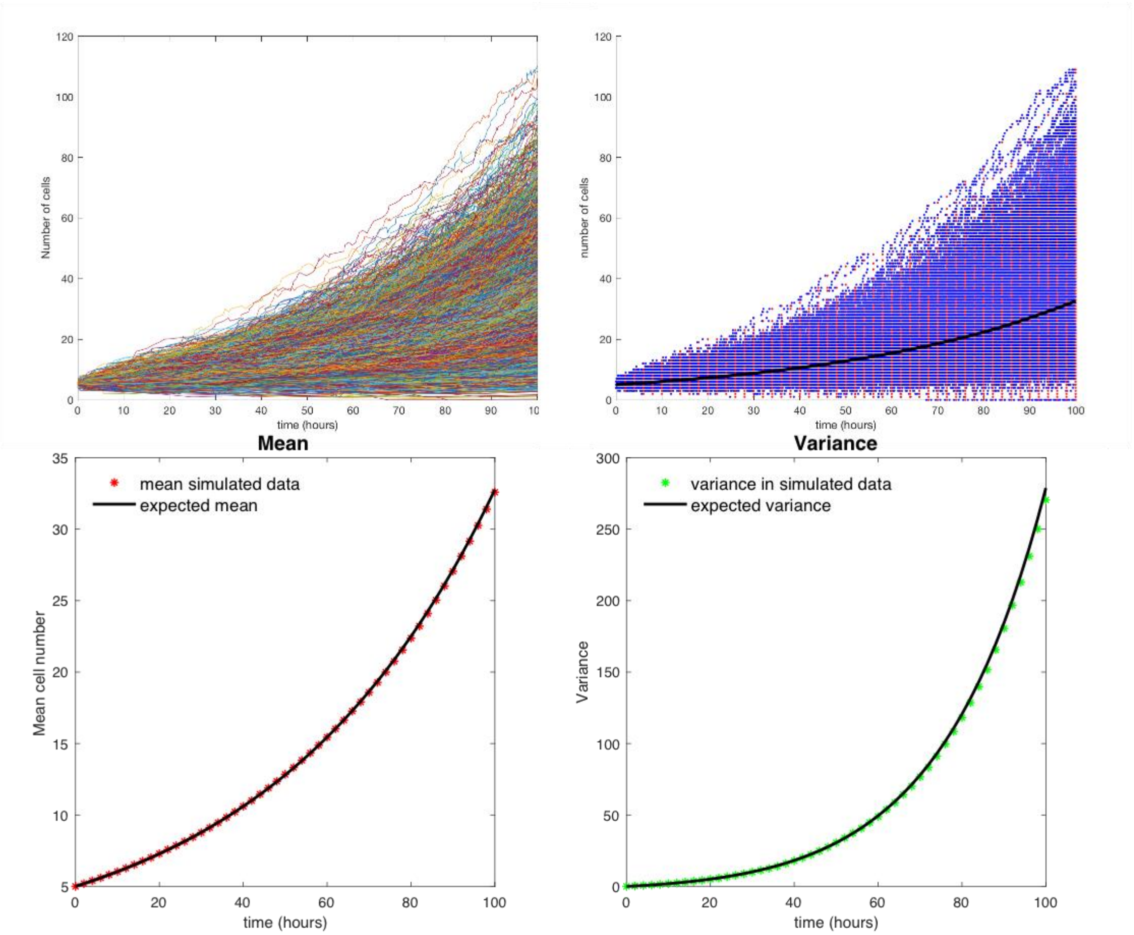


**Figure S2. Confirmation that moment approach derivations match measured mean and variance from simulated stochastic trajectories** (a) Example of stochastic growth model output from 5000 simulated cell number trajectories starting at a single cell with birth rate of b = 0.0238 and a death rate of d = 0.005, revealing the expected variability in growth dynamics that is not averaged out at low initial numbers (b) Stochastic growth trajectories uniformly samples every 4 hours (c) Measured mean at each time interval from simulated data with model expected mean as a function of time for the true parameters overlaid. (d) Measured variance at each time interval from simulated data with model expected variance as a function of time for the true parameters overlaid.

We repeated this procedure of simulating 5000 trajectories and computed the measured mean and variance for the remaining the strong Allee stochastic model family and the extended Allee model. We directly compared this to the solution of the moment approach approximation system of ODEs presented in Table 1.

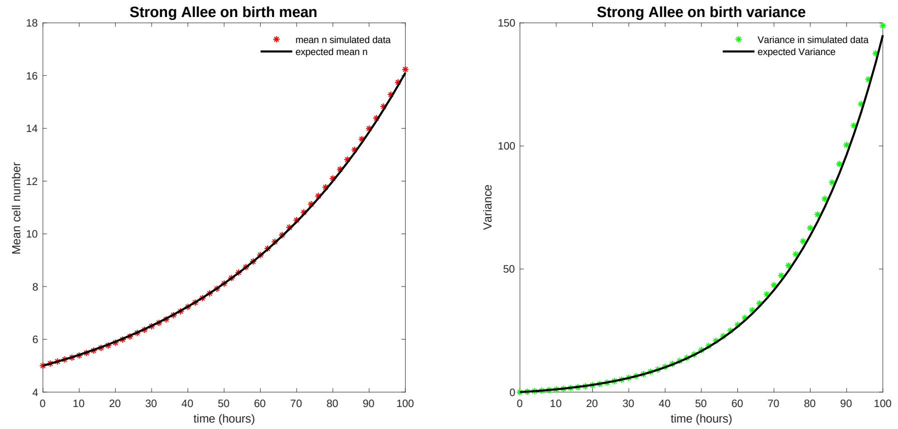


**Figure S3. Confirmation that moment approach derivations match measured mean and variance from simulated stochastic trajectories for strong Allee model on birth** (a) Measured mean at each time interval from simulated data with model expected mean as a function of time for the true parameters overlaid. (b) Measured variance at each time interval from simulated data with model expected variance as a function of time for the true parameters overlaid.

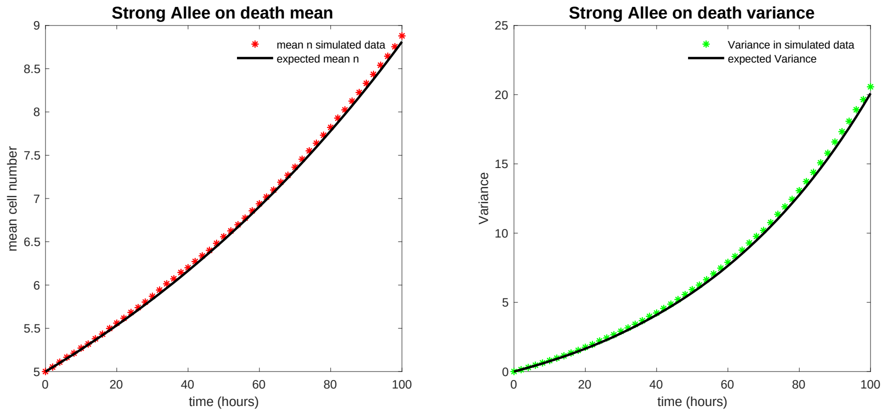


**Figure S4. Confirmation that moment approach derivations match measured mean and variance from simulated stochastic trajectories for strong Allee model on death** (a) Measured mean at each time interval from simulated data with model expected mean as a function of time for the true parameters overlaid. (b) Measured variance at each time interval from simulated data with model expected variance as a function of time for the true parameters overlaid.

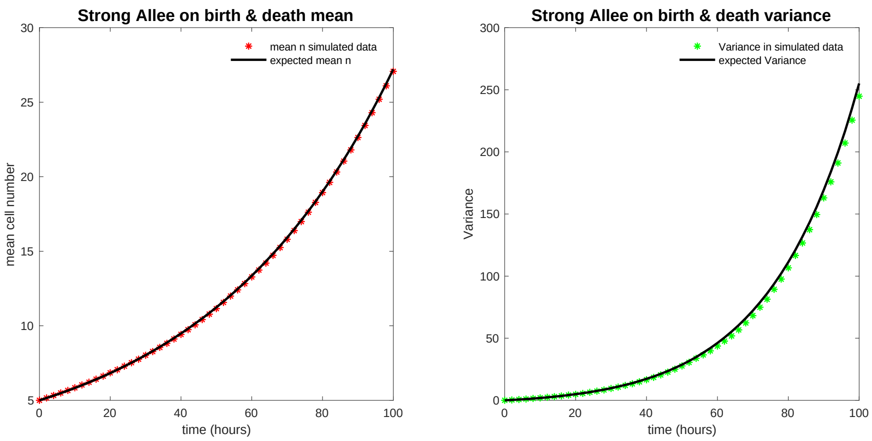


**Figure S5. Confirmation that moment approach derivations match measured mean and variance from simulated stochastic trajectories for strong Allee model on birth & death** (a) Measured mean at each time interval from simulated data with model expected mean as a function of time for the true parameters overlaid. (b) Measured variance at each time interval from simulated data with model expected variance as a function of time for the true parameters overlaid.

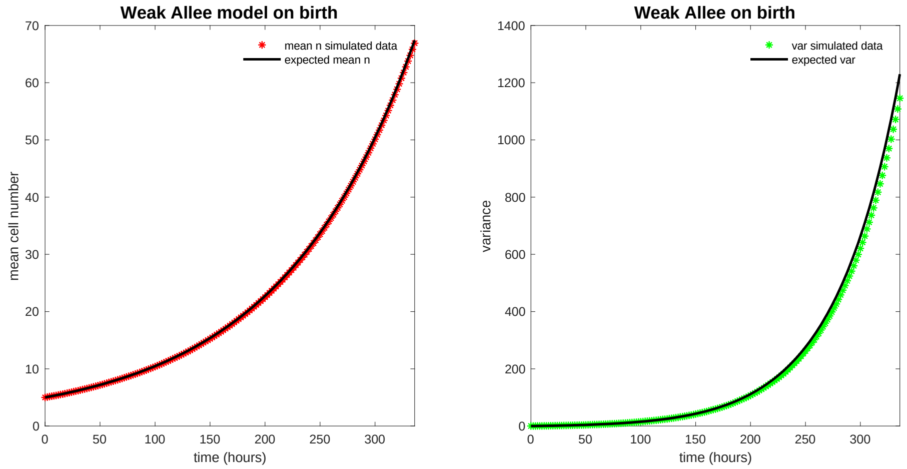


**Figure S6. Confirmation that moment approach derivations match measured mean and variance from simulated stochastic trajectories for weak Allee model on birth** (a) Measured mean at each time interval from simulated data with model expected mean as a function of time for the true parameters overlaid. (b) Measured variance at each time interval from simulated data with model expected variance as a function of time for the true parameters overlaid.

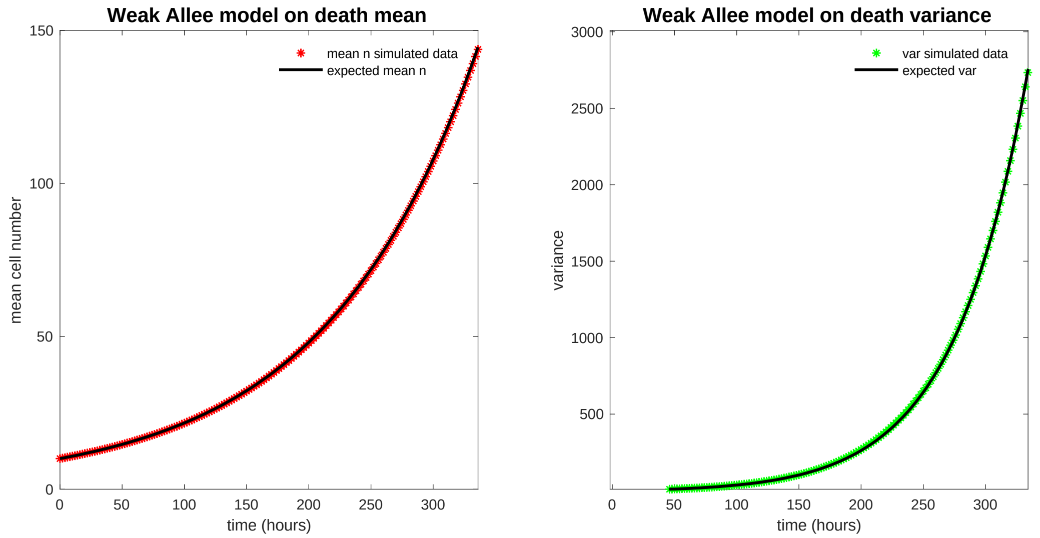


**Figure S7. Confirmation that moment approach derivations match measured mean and variance from simulated stochastic trajectories for weak Allee model on death** (a) Measured mean at each time interval from simulated data with model expected mean as a function of time for the true parameters overlaid. (b) Measured variance at each time interval from simulated data with model expected variance as a function of time for the true parameters overlaid.

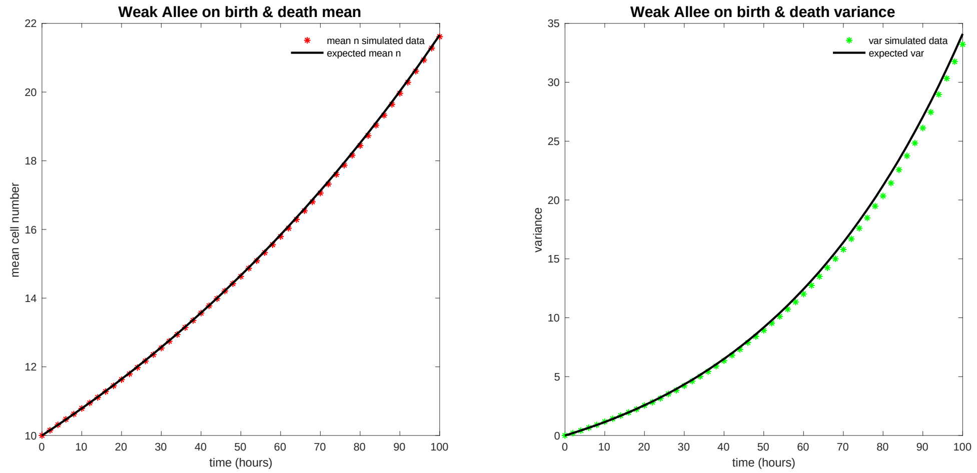


**Figure S8. Confirmation that moment approach derivations match measured mean and variance from simulated stochastic trajectories for weak Allee model on birth & death** (a) Measured mean at each time interval from simulated data with model expected mean as a function of time for the true parameters overlaid. (b) Measured variance at each time interval from simulated data with model expected variance as a function of time for the true parameters overlaid.

#### S4. Theoretical identifiability of the structural models using the differential algebra approach applied to the simple birth-death model as an example

The differential algebra approach(4)(5) requires equations be written to describe the process being modeled and the measurements available. In this case, we are modeling the linear two compartment system of differential equations that describes the time evolution of the mean and the variance in the bulk cell number. Because our experimental system is able to capture a high number of growth trajectories at low initial cell densities, we are able to measure the mean and variance throughout time for each initial condition. For the birth and death model, this leads to the following set of model and measurement equations from Table 1.

$$\dot{n} = (b - d)n$$

$$\dot{V} = 2(b - d)V + (b + d)n$$

$$y_1 = n$$

$$y_2 = V$$

Next, we can rewrite these in terms of the measurable outputs y1 and y2. We solve for n from the derivative of V differentiate, then set this equal to the derivative of n.

$$y_1 = \frac{\dot{y}_2 - 2(b - d)y_2}{b + d}$$

$$\dot{y}_1 = \frac{\ddot{y}_2}{b + d} - \frac{2(b - d)\dot{y}_2}{b + d} = (b - d)y_1$$

$$0 = \frac{\ddot{y}_2}{b + d} - \frac{2(b - d)\dot{y}_2}{b + d} - (b - d)y_1$$

The coefficients in front of the measurable outputs are the identifiable parameter combinations, which we set equal to $a_1$, $a_2$, and $a_3$. We then use substitution and replacement to solve for the parameters b & d in terms of the identifiable combinations.

$$a_1 = \frac{1}{b + d}$$

$$a_2 = \frac{-2(b - d)}{b + d}$$

$$a_3 = b - d$$

$$b + d = \frac{1}{a_1}$$

$$b = \frac{1}{a_1} - d$$

$$a_2 = \frac{2(\frac{1}{a_1} - 2d)}{\frac{1}{a_1}}$$

$$d = \frac{2 - a_2}{4}$$

$$b = \frac{1}{a_1} + \frac{a_2}{4} - \frac{1}{2}$$

If we can isolate each parameter in terms of identifiable combinations (a1, a2, and a3) alone, then the parameters are structurally identifiable, as is shown here by the isolation of b & d. Note that the unique identifiability of b and d, not just (b-d) would not be identifiable without the measurement of the variance (specifically because the time derivative of the variance is proportional to b+d), as is explained in Figure S1. We performed this analysis for all seven stochastic model structures, and found all parameters to be uniquely structurally identifiable.

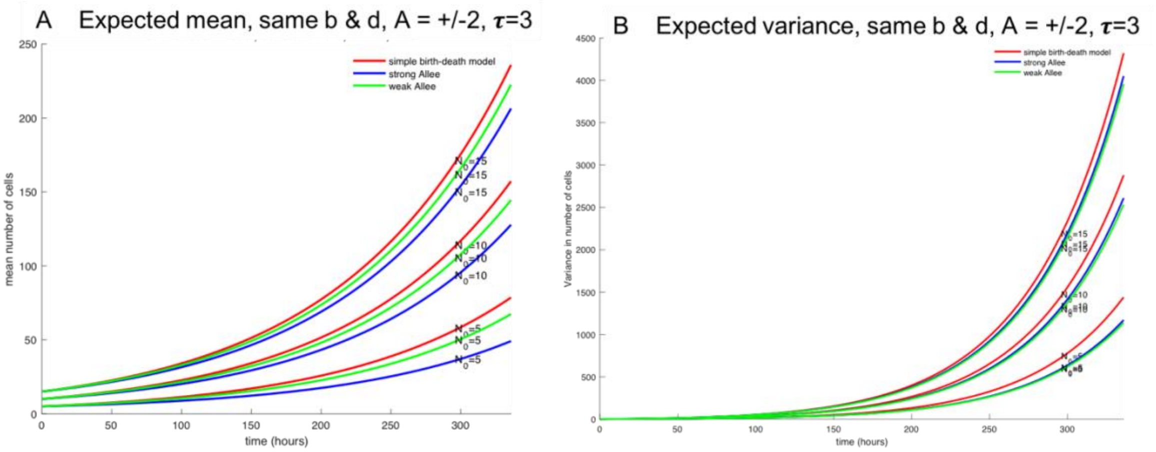


**Figure S9. Comparison of similar simple birth-death, strong, and weak Allee expectations for the time evolution of the mean and variance**. (a) Expected time evolution of the mean cell number for the simple birth-death model (red), the strong Allee model on birth (blue) and the weak Allee model on birth (green) with the same birth and death rates for all but with A=2 for the strong Allee model and A=-2, $\tau$=3 for the weak Allee model indicates significant differences in trajectories for $N_0$ = 5, 10, & 15 (b) Expected time evolution of the variance in cell number for the same initial conditions and parameters.

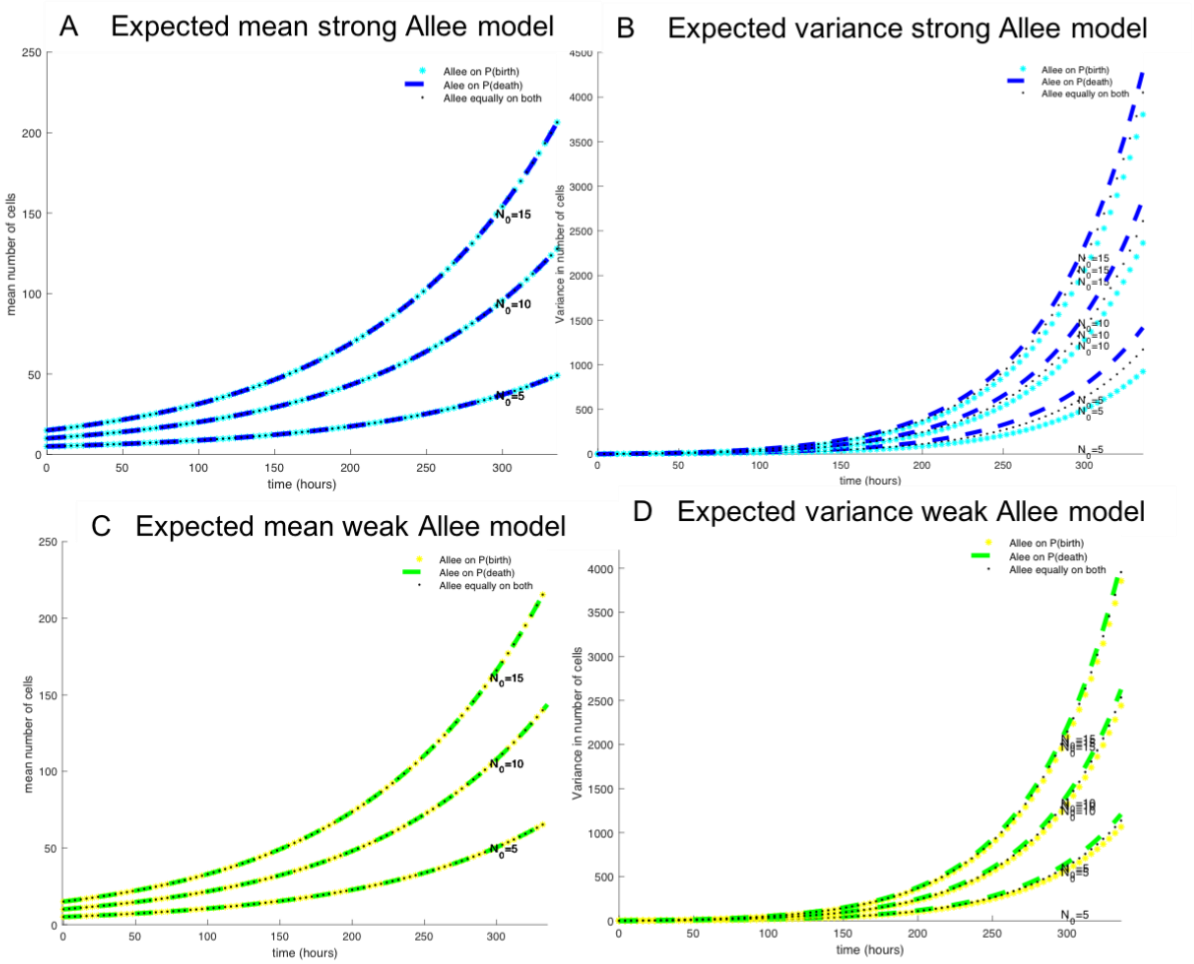


**Figure S10. Demonstration of effect of Allee mechanism on birth or death probability on the variance.** (a) As expected, for constant parameters the mean cell number in time for the strong model is the same for the strong Allee model on birth, death, or both. (b) The expected time evolution of the variance for the strong Allee model on the birth probability (cyan), death probability (dark blue) and them equally (black) (c) As expected, for constant parameters the mean cell number in time for the weak Allee model is the same for the strong model on birth, death, or both. (b) The expected time evolution of the variance for the strong model on the birth probability (yellow), death probability (green) and them equally (black).

#### S5. Model Selection using Bayesian Information Criterion and BIC weights

To investigate competing hypotheses about the underlying structure of tumor growth dynamics, the seven distinct stochastic models were compared using the Bayesian information criterion (BIC) (6) (7). The BIC takes into account both goodness of fit of the model and penalizes for complexity of the model in terms of number of parameters, and has been shown to be an inexpensive approximation to the Bayes factors, which gives the favor of a model over another (7). In order to ensure the method was not overweighing goodness of fit, the data was down-sampled from the true data collection interval of every 4 hours to every 36 hours to demonstrate that down-sampling changed the magnitudes of the BIC values but did not affect the order of the BIC values of each model relative to one another (Figure S11). To evaluate statistical significance between models with BIC values that were very close to one another, the methods presented in Waenmakers & Farrell et al (8) of BIC weighting were used which are given by:

$$w_i(BIC) = \frac{\exp(-\frac{1}{2}\Delta_i(BIC))}{\sum_{k=1}^{K}\exp(-\frac{1}{2}\Delta_k(BIC))}$$

From this equation, each model is assigned a relative weight, whose sum add to 1 based on the probability that it is the most parsimonious model to describe the data.

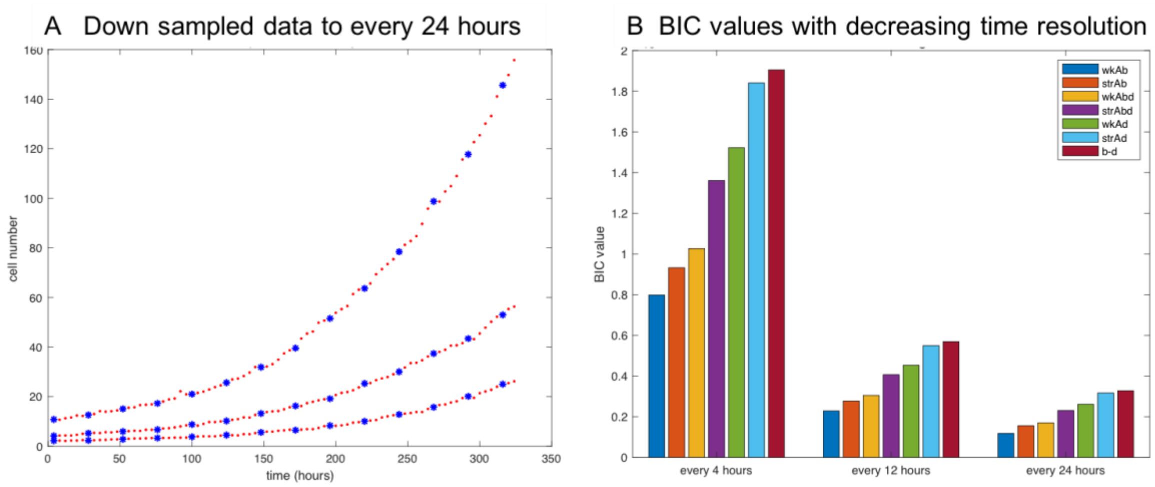


**Figure S11. Decreased time resolution of data does not change fitting results, weak Allee model on birth is consistently chosen.** (a) Example of down sampled time resolution from original data (red) to data every 24 hours (blue) (b) BIC values for each model fit at data sampled every 4, 12, and 24 hours respectively reveals weak Allee model has consistently the lowest BIC value and is chosen every time.

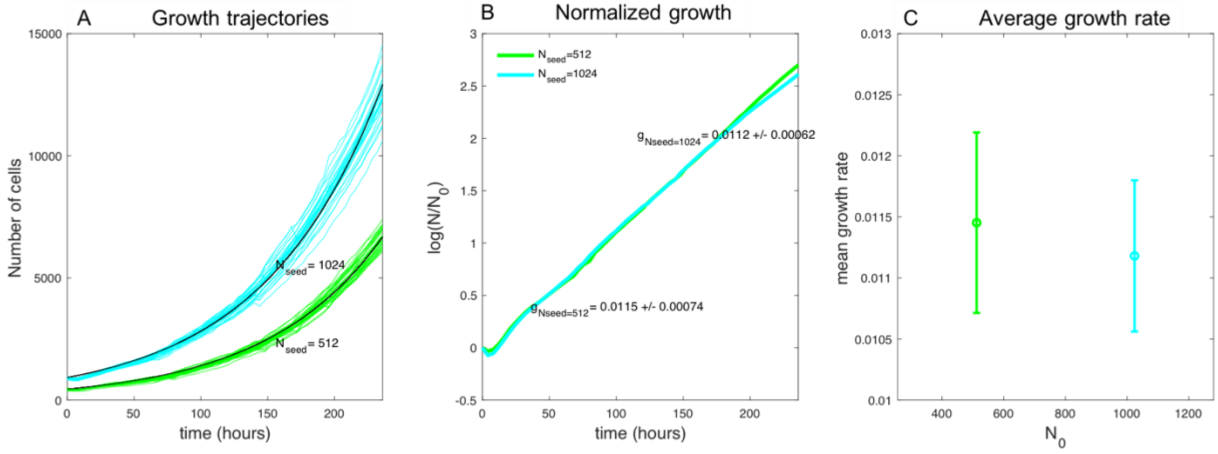

**Figure S12. Normal cell culture density exhibits expected constant growth rate** (a) Thirty growth rate trajectories for seeding of $N_0$ = 512 (green) & $N_0$ = 1024 (cyan) (b) Normalized cell number in time by $N_0$ reveals expected constant growth rate (c) Average growth rate and for $N_0$=512 (green) of g = 0.0112+/- 0.00062 and $N_0$ = 1024 g = 0.0115 +/- 0.00074

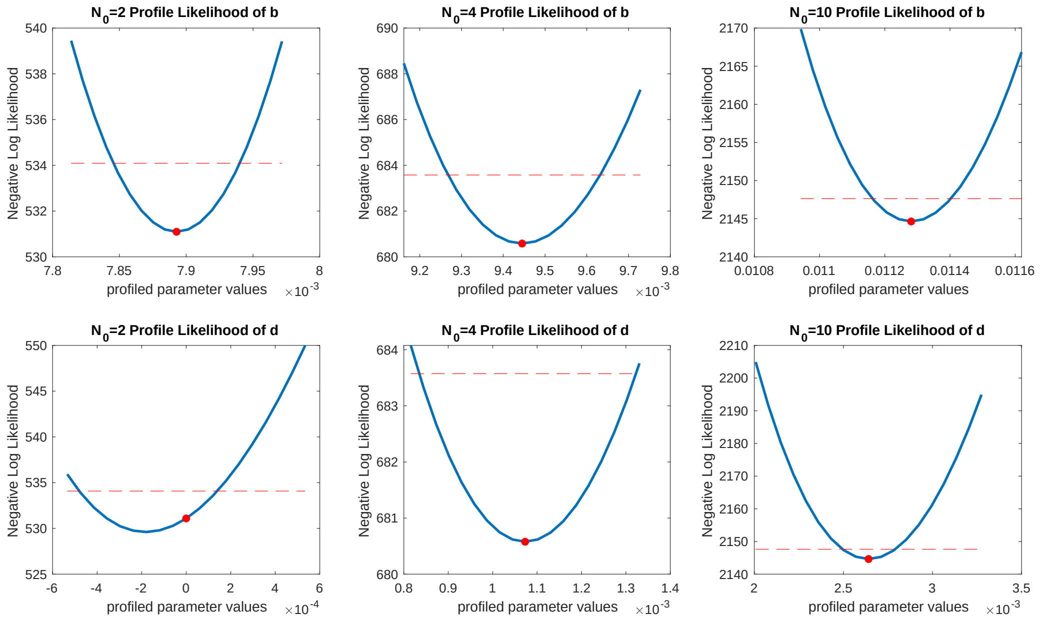


**Figure S13: Profile likelihood analysis on birth and death rates for individual cell numbers $N_0$= 2, 4, & 10 reveals practical identifiability of the birth and death rate parameters for datasets of each individual group of $N_0$ trajectories.** (a), (b), (c). Profile likelihood analysis on birth rate parameter for $N_0$=2, 4, &10 respectively. (d), (e), (f). Profile likelihood analysis on death rate parameter for $N_0$ = 2, 4, & 10 respectively.

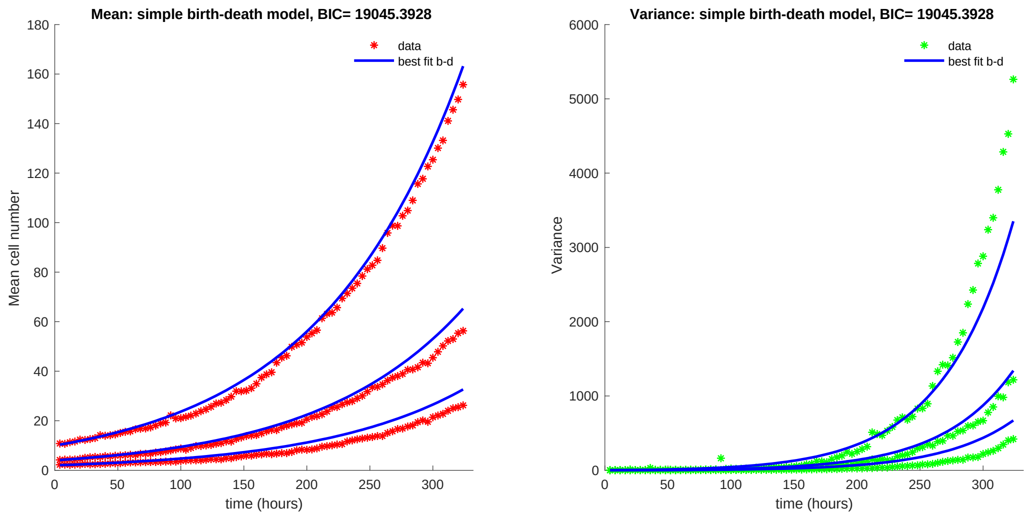


**Figure S14: Data fit to birth-death model results in a BIC= 1.9e4**. (a) Mean of the data (red) to the best fitting birth-death model mean (blue). (b) Variance of the data (green) to the best fitting birth-death model variance (blue).

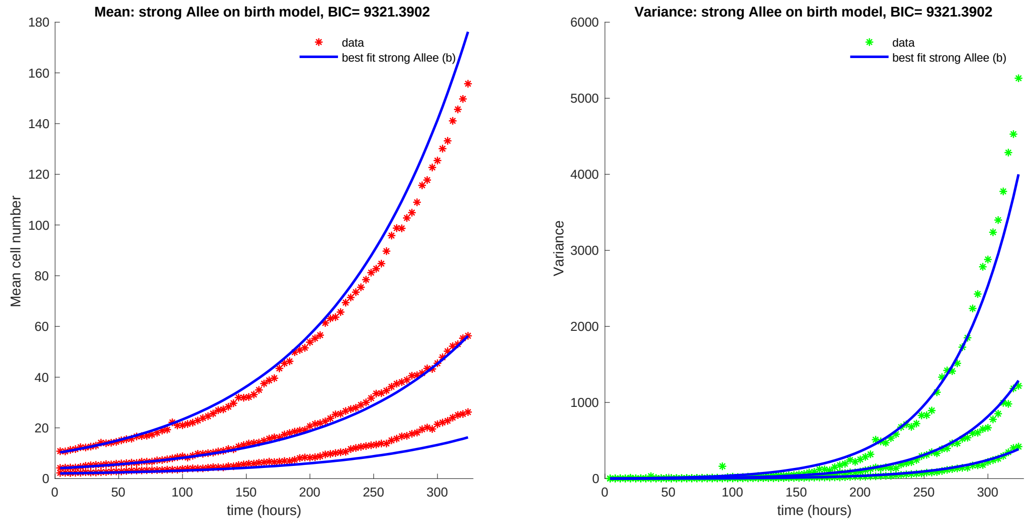


**Figure S15: Data fit to strong Allee on birth model results in a BIC= 9e3**. (a) Mean of the data (red) to the best fitting strong Allee on birth model mean (blue). (b) Variance of the data (green) to the best fitting strong Allee on birth model variance (blue).

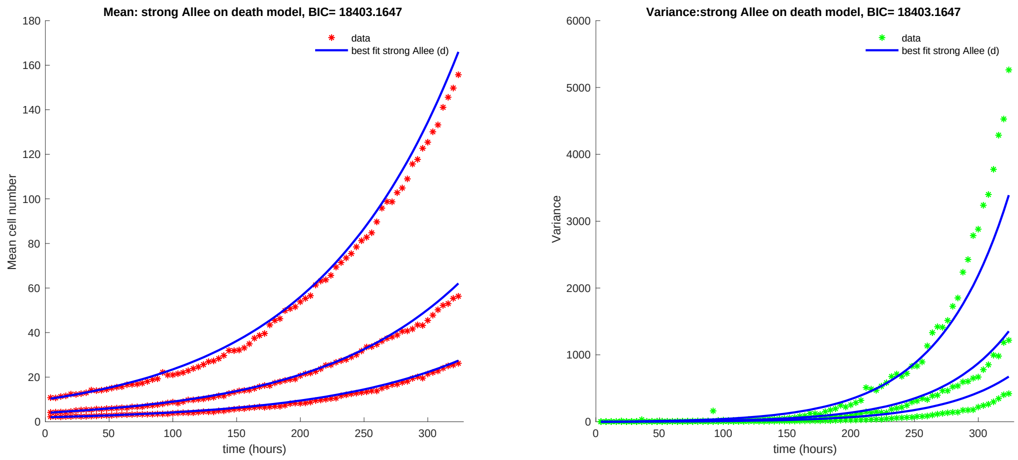


**Figure S16: Data fit to strong Allee on death model results in a BIC= 1.8e4**. (a) Mean of the data (red) to the best fitting strong Allee on death model mean (blue). (b) Variance of the data (green) to the best fitting strong Allee on death model variance (blue).

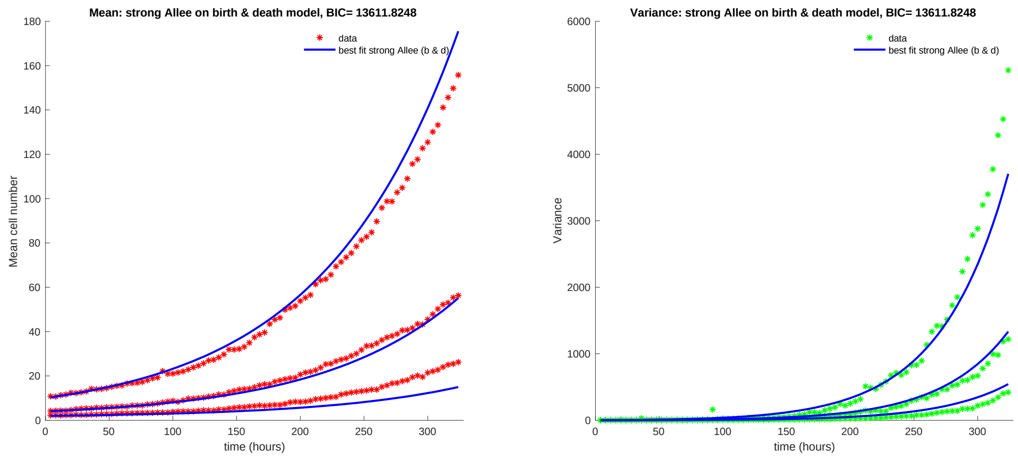


**Figure S17: Data fit to strong Allee on birth & death model results in a BIC= 1.4e4**. (a) Mean of the data (red) to the best fitting strong Allee on birth & death model mean (blue). (b) Variance of the data (green) to the best fitting strong Allee on birth & death model variance (blue).

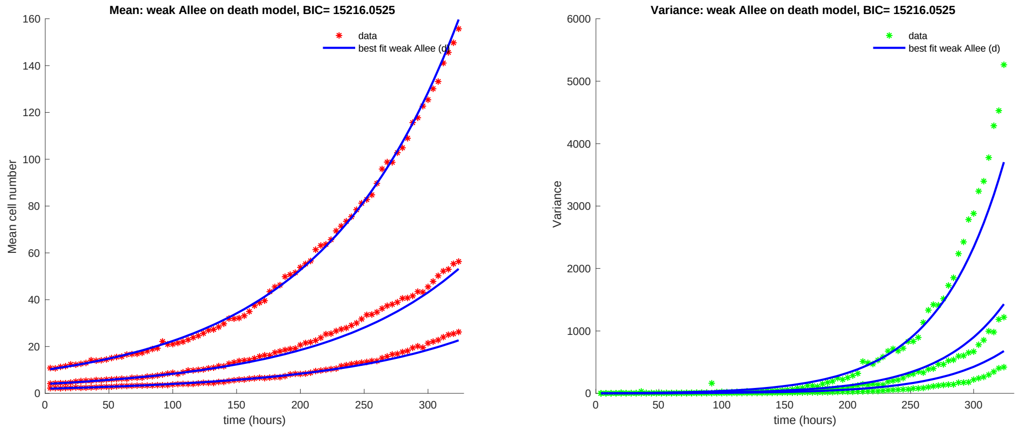


**Figure S18: Data fit to weak Allee on death model results in a BIC= 1.5e4**. (a) Mean of the data (red) to the best fitting weak Allee on death model mean (blue). (b) Variance of the data (green) to the best fitting weak Allee on death model variance (blue).

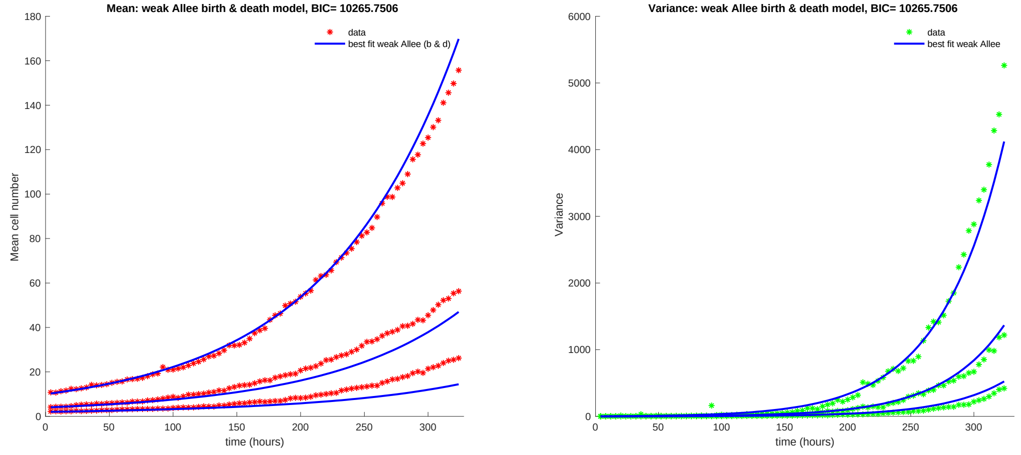


**Figure S19: Data fit to weak Allee on birth & death model results in a BIC= 1.0e4**. (a) Mean of the data (red) to the best fitting weak Allee on birth & death model mean (blue). (b) Variance of the data (green) to the best fitting weak Allee on birth & death model variance (blue).

### References

1. Gillesple DT. Exact Stochastic Simulation of Coupled Chemical Reactions. J Phys Chem. 1977;81(25):2340–61.
2. Gillespie DT. The chemical Langevin equation. J Chem Phys. 2014;297(2000).
3. Houchmandzadeh B. Extracting moments from Master Equations. ArXiv. 2009;1(2):1–14.
4. Meshkat N, Sullivant S, Eisenberg M. Identifiability Results for Several Classes of Linear Compartment Models. Bull Math Biol. 2015;77(8):1620–51.
5. Brouwer AF, Meza R, Eisenberg MC, Arbor A. A systematic approach to determining the identifiability of multistage carcinogenesis models. Risk Anal. 2018;37(7):1375–87.
6. Raftery A. Bayes Factors and BIC. Sociol Methods Res. 1999;27(3):411–27.

7. Loos C, Moeller K, Fröhlich F, Hucho T, Hasenauer J. A Hierarchical, Data-Driven Approach to Modeling Single-Cell Populations Predicts Latent Causes of Cell-To-Cell Variability. Cell Syst. 2018;6(5):593–603.e13.
8. Wagenmakers E, Farrell S. AIC model selection using Akaike weights. Psychon Bull Rev. 2004;11(1):192–6.
